# Uncoupling of seagrass host selection and succession for microbial guilds in meadow chronosequence

**DOI:** 10.64898/2026.03.24.714081

**Authors:** Prasha Maithani, Clarence Wei Hung Sim, Sujatha Srinivas, Zi Cen Kwek, Rebecca Josephine Case

## Abstract

Succession is an ecosystem building process in which a habitat and its community interact predictably by increasing diversity, habitat engineering, and ultimately reaching a climax community, where other ecological processes influence its dynamic. Key to succession is the establishment of primary producing habitat forming species, which drives niche differentiation leading to increasing diversity. Here, we use the primary colonizing and habitat forming seagrass, *Halophila ovalis*, to demonstrate that it drives bacterial succession in a meadow ecosystem, and its microbiome, both rhizoplane and phylloplane, are under host selection. Many of the characteristics attributed to plants for habitat modification are microbial processes such as nitrogen fixation and sulfide detoxification and succession is often extrapolated to such processes. To determine if succession (increasing diversity) or selection (reducing diversity) drives changes in diversity (16S rRNA gene) or habitat modifying processes (*nifH*, *soxB*, *aprA*, *dsrA*), molecular analysis was performed along chronosequences (as a proxy for succession) of seagrass patches. Bacterial communities were sampled within the meadow ecosystem and the microbiomes of *H. ovalis* (both rhizoplane and phylloplane). Genes involved in biogeochemical cycling are differentially impacted within the microbiome and meadow sediments, with only *nifH* under succession. All genes from all niches sampled for community analysis are under directional community trajectories, despite being subjected to distinct ecological processes, signifying that many ecological processes, including succession and host association, drive community assemblage.

## INTRODUCTION

Ecological succession encompasses the directional and predictable changes in biological communities that occur following primary colonisation of new habitats or disturbance of existing habitats. Early colonists, or pioneer taxa, are known to modify local habitat conditions through physical, chemical, and biological feedbacks that can either facilitate or inhibit the establishment of later-arriving species (Connell & Slatyer, 1977). Key to successionally driven processes are habitat forming species that support increasing diversity within the ecosystem by providing habitat and as they are characteristically phototrophs (plants, grasses, algae, corals), they also feed the ecosystem through photosynthesis. The ecosystem’s increasing diversity is driven by shifts in species composition, resource availability, and biotic interactions (Connell & Slatyer, 1977; Odum, 1969). Early succession models are depicted as the changing assemblage of habitat forming species and often define the ecosystem once reaching a climax community (e.g. forests, meadows, coral reefs). These two distinct phases of succession: (1) establishment of an ecosystem and (2) a dynamic climax community, are not strictly discrete. However, diversity should predictably increase along a gradient of time during establishment and be dynamic, perhaps entering alternate states, within a climax community. Although originally developed in the context of terrestrial plant communities, succession theory is broadly applicable to any habitat-forming organism capable of engineering its environment (Ortiz-Alvarez et al., 2018).

Seagrasses and the meadows they form represent such habitat-forming or foundation species. As primary producers, seagrasses are the foundation of coastal food webs and support diverse faunal communities (Lamb et al., 2017; Thomson et al., 2015). In addition, the holobiont metabolic activity and physical structure exert strong engineering effects: seagrass holobionts modify sediment redox gradients, stabilise sediment, attenuate hydrodynamic energy, and regulate nitrogen, sulfur, and carbon cycling in coastal waters and sediments (Duarte et al., 2010; Mazarrasa et al., 2015; Ward et al., 2022). In doing so, they create physicochemical niches that structure the seagrass microbiome (Simon et al., 2019; Tarquinio et al., 2019), fostering host-selected nitrogen and sulfur cycling bacteria uniquely adapted to the conditions the plant itself generates (Ugarelli et al., 2017). However, such interactions are not unidirectional. Indeed, many functional attributes of seagrasses are mediated by their microbiome as bacteria and archaea have greater metabolic versatility than eukaryotes and are key players in carbon, nitrogen and sulfur biogeochemical cycling. And while host species, such as seagrasses, can place selective pressure on their microbiome, succession theory predicts they would drive increasing diversity within the ecosystem (i.e. sediments) during establishment, a hypothesis largely untested in seagrass meadows.

Plant-associated microbiomes are strongly structured by contrasts between above-and below-ground tissues. In seagrasses, the phylloplane comprising leaf and stem tissues is oxygenated, exposed to sunlight, intertidally submerged in seawater and exposed to the atmosphere. Enriched in sugars, the phylloplane fuels heterotrophic bacteria recruitment from the water column, ultimately forming a biofilm on the leaf surface (Crump et al., 2018; Ugarelli et al., 2017; Vogel et al., 2020). This biofilm is indistinguishable in composition on *Zostera marina* from the surrounding water (A. K. Fahimipour et al., 2017). However, when the surface water film of motile and reversibly attached microbes is removed, a distinct phylloplane community was identified (Iqbal et al., 2022), demonstrating that seagrass leaves can exert selective pressure on the leaf biofilm community.

The rhizosphere, comprising roots and rhizomes embedded in seawater saturated sediment, forms a chemically heterogeneous microenvironment shaped primarily by root oxygen release at the growing tip and high sucrose accumulation (Hennion et al., 2019; Sogin et al., 2022), resulting in structured microbial communities, and strong pH, oxygen, redox and biogeochemical gradients that fluctuate diurnally with host photosynthetic activity. The reversibly attached and motile microbial community is able to respond to the diurnally fluctuating gradients (Banister et al., 2022; Brodersen et al., 2024; Brodersen et al., 2018). Within the rhizosphere, the rhizoplane refers specifically to the root-surface biofilm, a host-selected and tightly adherent microbial assemblage that is compositionally distinct from the more dynamic surrounding rhizosphere (Barea et al., 2005). Rhizoplane-associated bacteria often exhibit specialised adaptations for host association, including exopolysaccharide production by taxa such as *Azotobacter*, which enhances plant growth and stress tolerance (Gauri et al., 2012), and the capacity of symbiotic rhizobacteria to induce structural and physiological modifications in plant roots leading to the formation of specialised nodular structures (Aasfar et al., 2021).

Despite compositional differences, both the phylloplane and rhizoplane communities contribute to nutrient cycling and plant health. Nitrogen-fixing bacteria and archaea that provide biologically available nitrogen essential for host biomass production inhabit both tissues (Crump et al., 2018; Mohr et al., 2021). In marine sediments, anaerobic conditions drive sulfate reduction, resulting in the accumulation of hydrogen sulfide toxic to eukaryotic cells (Hasler-Sheetal & Holmer, 2015; Jiang et al., 2016). Consequently, the seagrass delivers oxygen to the growing root tip (Martin et al., 2019) and the sulfur-cycling consortia that are essential for mitigating sulfide toxicity in eukaryotic cells occur predominantly in the rhizosphere and are conserved across diverse seagrass species and regions (Ashkaan K. Fahimipour et al., 2017; Hasler-Sheetal & Holmer, 2015; Rabbani et al., 2021).

Succession in seagrass ecosystems mirrors classic terrestrial plant succession theory but occurs at the scale of vegetated patches, which expand into unvegetated sediment, eventuating in patches merging into a meadow ecosystem. Seagrasses spread horizontally through clonal rhizome extension (Tomlinson, 1974), giving rise to new shoots that serve as nascent ecosystems for microbial colonisation (Jensen et al., 2007). Beyond initial host colonisation, growing shoots can drive microbial succession over time as increasing primary productivity increases organic carbon and oxygen in sediments, the associated microbiome drives biogeochemical cycling that alters the redox chemistry of sediments, and seagrasses also modify physical properties through wave attenuation. These early patches undergo shifts in species abundance, functional potential, and microbial interactions as they mature (García-Martínez et al., 2005; Martin et al., 2018). Microbial succession has been documented across multiple marine systems, including microalgal microbiomes (Majzoub et al., 2019; Mönnich et al., 2020), macroalgal microbiomes (Longford et al., 2019), lichen bacterial communities (Mushegian et al., 2011), hydrothermal vents (Hou et al., 2020), and sinking particles (Stephens et al., 2024), demonstrating that host-associated microbial communities restructure predictably over time, and timescales for microbial assemblage are much shorter than microbe succession. In seagrasses, their colonial expansion and known growth rates in patches enable a chronosequence to predictably infer their relative age between adjacent plants, enabling inference of successional changes (Friess et al., 2023; Walker et al., 2010).

Singapore offers a valuable model for studying early-stage seagrass succession. Despite severe coastal urbanisation and an estimated 45% habitat loss since the 1970s (Yaakub et al., 2014), sites like Chek Jawa on Palau Ubin remain protected and host exceptionally high tropical seagrass diversity (Fortes et al., 2018). With the site’s environmental stability and discrete patch-based meadows, Chek Jawa provides an ideal setting for investigating microbiome assembly across natural successional gradients. Here, we investigate how bacterial and archaeal communities assemble during early stages of seagrass patch development. Specifically, we aim to: (1) determine whether the host-associated microbiome (rhizoplane and phylloplane) and the sediment microbiome exhibit similar successional patterns; (2) test whether functional diversity scales with organismal (taxonomic) diversity across successional stages; (3) compare the influence of succession between community and functional assemblages.

## MATERIALS AND METHODS

### Study site, chronosequence and sampling

All samples were collected at Chek Jawa Wetlands (1.409075 N, 103.992737 E), on Pulau Ubin in northern Singapore (Fig. 1A). *Halophila ovalis* and sediment (Fig. 1B-D) were sampled from the intertidal zone during a low tide of <0.4 m relative to chart datum, when plants are not submerged in coastal water but are exposed to the air. Chek Jawa is a natural meadow with limited access to the public and has not undergone reclamation (McKenzie et al., 2016; Yaakub et al., 2014). *H. ovalis* is the dominant seagrass species, with *Cymodocea rotundata* and *Halodule uninervis* co-occurring in patches. *Halophila major* and *Halophila minor* were excluded from *H. ovalis* sampling (Kwan et al., 2023). Surface oxic sediment (∼ 1 cm depth) in the meadow is sand-coloured and covers grey-black anoxic sediments (Fig. 1D) that extend below seagrass roots. All sediments collected were grey-black, indicative that they were anoxic.

**Figure 1.**
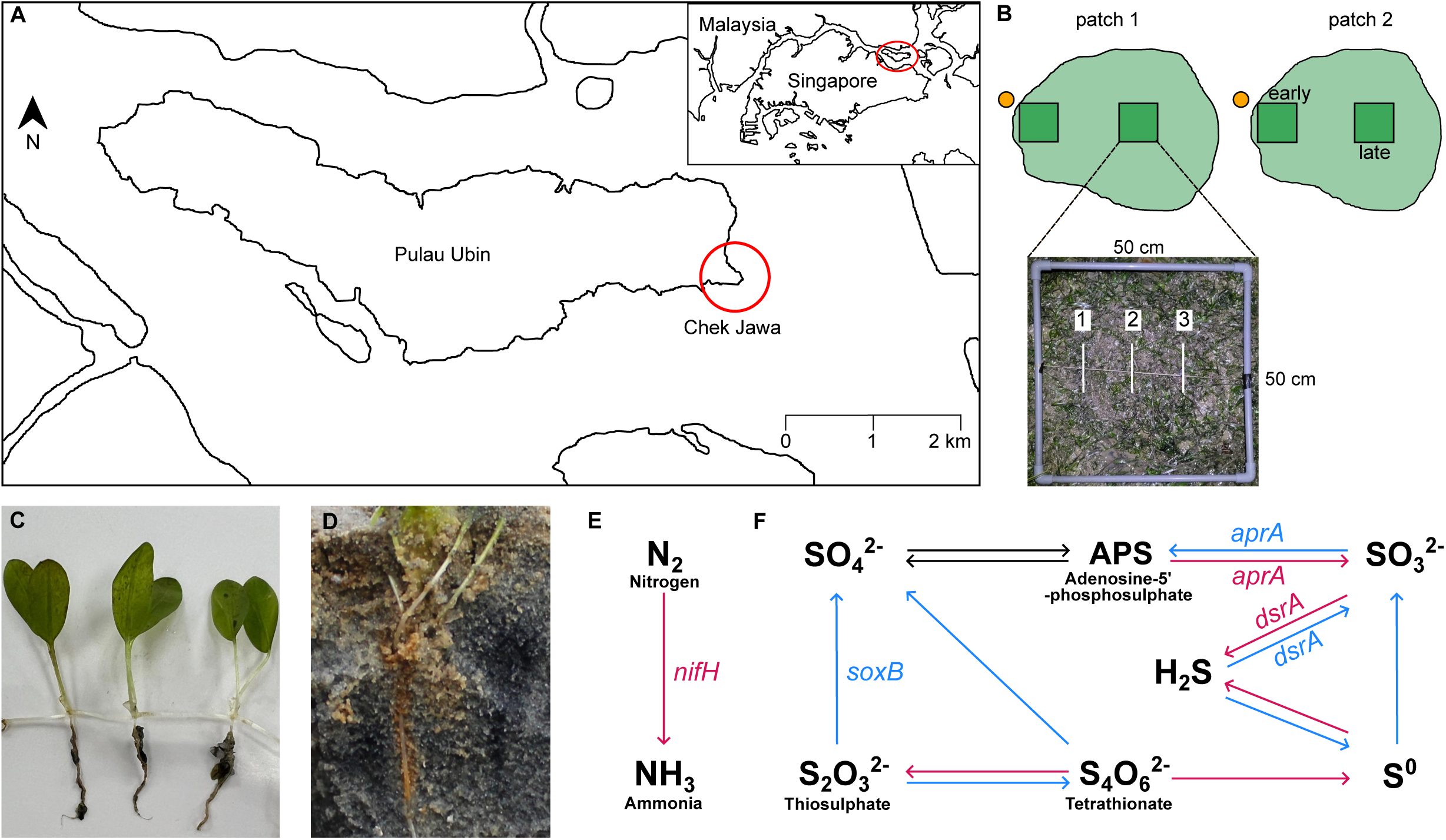
Seagrass, sediments, chronosequence sampling location and study design. **(A)** Map of the study site showing Chek Jawa (red circle), Pulau Ubin and Singapore (inset). A seagrass patch is the unit from which a meadow forms so a **(B)** chronosequence was sampled from two patches (light green), with small scale spatial sampling from three equidistant positions (1, 2, 3) within 50 cm X 50 cm quadrats, large scale spatial sampling of early and later successional communities between quadrats (dark green squares), and pre-succession sediment outside the patch (orange circles). Sediment was sampled within 2 cm of the root but not attached to the plants. **(C)** Three *Halophila ovalis* plants along its rhizome, showing above and below ground organs **(D)** Sediment core from Chek Jawa, showing oxic (sand coloured) and anoxic (grey-black coloured) sediment layers. Orange (iron oxide) colouration on the root and the extending gradient from the roots of sand togrey-black (iron sulfide) sediment visualises the chemical gradient in the rhizosphere. **(E)** The nitrogenase iron protein (*nifH* gene) is part of the nitrogenase complex that reduces nitrogen to ammonia during nitrogen fixation. **(F)** The sulphate thiohydrolase (*soxB* gene) is a di-manganese containing protein of the sulphur oxidation (Sox) complex that oxidises thiosulphate to sulphate. The adenosine-5’-phosphosulphate reductase (*aprA* gene) catalyses the reversible reduction of adenosine-5’-phosphosulphate to sulphite and the dissimilatory sulphite reductase (*dsrA* gene) catalyses reduction of sulphite to sulphide. The *aprA* and *dsrA* genes can function in both dissimilatory sulphate reduction and sulphur oxidation pathways, depending on redox conditions. Red and blue arrows represent reductive and oxidative reactions, respectively.

Sampling was done within patches (Fig. 1B) at the Chek Jawa meadow (Fig. 1A), where a patch is defined as a single unit of the seagrass meadow ecosystem, and a continuous area of seagrass cover at least 4 m maximum width, bordered by unvegetated sediment on at least one side. Two patches were sampled as chronosequences at small (within quadrat) and larger (between quadrats) spatial scales (Fig. 1B), to capture early and later succession during patch expansion, but before patches weren’t confluent with the meadow. A 50 x 50 cm quadrat was placed at the expanding edge of patch, and another inside the patch, with a minimal 3 m distance between the quadrats (Fig. 1B). In each quadrat, three equidistant *H. ovalis* plants free of epiphytes were selected, and 1 cm^3^ of sediment was collected that was not attached or in direct contact with the plant, but within 2 cm of the plant root, at a depth of 4 cm (ensuring sediments were grey-black), using 10 cm^3^ sterile syringes with modified tips. Additionally, unvegetated sediment was collected outside each patch, at the same depth and with the same appearance. Plants were collected using a sterilized shovel and were gently agitated in ambient seawater and subsequently rinsed with filter-sterilized, autoclaved seawater to remove reversibly attached sediments and organisms, and to ensure that we have the attached, or biofilm microbiome. The plants were immediately sectioned into above-ground (phylloplane, comprising the stem and leaves) and below-ground (rhizoplane comprising the rhizome and roots) organs (Fig. 1C) using sterile blades. All samples were immersed in RNAlater, and placed on ice in the field, then within 3 h stored at 4 °C overnight, then at -20 °C until further processing.

### DNA extraction, PCR and sequencing

Environmental DNA (eDNA) was extracted from *H. ovalis* and sediments using the DNeasy PowerSoil Pro Kit (Qiagen, Germany), following the manufacturer’s protocol, with additional homogenization performed using the FastPrep-24^TM^ 5G Instrument (MP Biomedicals, Singapore) for three cycles at 4 m/s for 30 s, with 30 s rest in between. Field samples were thawed on ice prior to extraction and a minimum of 60 mg per sample was used for extraction. All extracted DNA was quantified using Qubit Fluorometer and Qubit dsDNA BR (Broad Range) assay kit (ThermoFisher Scientific, USA), following the manufacturer’s protocol. Negative extraction controls included to assess contamination showed DNA concentrations < 0.1 ng/µL by Qubit and no detectable bands after PCR amplification.

To investigate successional changes in microbial communities, and in nitrogen- and sulphur-cycling taxa, gene fragments of the 16S rRNA gene, *nifH*, *soxB*, *aprA* and *dsrA* (Fig. 1E & F) were PCR amplified (Table S1). All primers were modified to include Illumina overhang adapter sequences, and recombinant bovine serum albumin (rBSA) was added to all PCR reactions. All reactions contained a minimum of 25 ng of DNA, and included an initial denaturation step at 94 °C for 180 s. The 16S rRNA gene fragment was amplified using the 515F/806R primers (Caporaso et al., 2011) as previously described (Yan et al., 2021) but with a reduction from 35 to 25 cycles of the following: 94 °C for 45 s, 50 °C for 60 s, and 72 °C for 90 s, with a final extension at 72 °C for 10 minutes. The polF/polR (Poly et al., 2001), 693F/1164R (Petri et al., 2001), aprA-1-FW/aprA-5-RV (Aoki et al., 2015), and DSR1F/DSR-R (Gao et al., 2022) primer pairs were used to amplify the *nifH*, *soxB*, *aprA* and *dsrA* genes, respectively. Cycling conditions for functional genes included 30 cycles of 94 °C for 30 s, annealing for 60 s and 72 °C for 60 s, with a final extension of 72 °C for 7 minutes. Annealing temperatures were determined based on the primer pairs and were as follows: 59 °C (*nifH*), 53 °C (*soxB*), 53 °C (*aprA*) and 59 °C (*dsrA*). PCR conditions were optimised to reduce the number of amplification cycles so to reduce artifacts, reduce the annealing temperature to increase the diversity amplified and the nested PCR protocol was removed as this amplifies PCR bias and limits interpretation of proportional abundance data (Polz & Cavanaugh, 1998; Thompson et al., 2002). Negative PCR controls included to assess contamination showed no detectable bands on gel after amplification.

All PCR products were visualised on a 1% Tris-acetate-EDTA (TAE) buffer agarose gel to check for single bands of the correct size, then purified using AMPure XP beads (Beckman Coulter, USA), following the manufacturer’s protocol. Purified DNA was quantified using the Qubit Fluorometer and the Qubit 1X dsDNA (HS) assay kit (ThermoFisher Scientific, USA), following the manufacturer’s protocol. A minimum amount of 50 ng of amplicon was submitted for sequencing on the Illumina MiSeq platform (Singapore Centre for Environmental Life Sciences Engineering), using V3 chemistry for 300 bp paired-end reads.

### Bioinformatic analysis

All sequencing reads were demultiplexed, and primers and adapters removed using *Cutadapt* version 4.0 (Martin, 2011). The *DADA2* package version 1.26 (Callahan et al., 2016) implemented in R version 4.2.2 was used to generate amplicon sequence variants (ASVs). Briefly, reads were trimmed and filtered with a maximum expected error of 2, error corrected, used to infer variants, merged, and cleared of chimeras. All subsequent analyses were performed using R version 4.5.1 (R Core Team, 2025).

16S rDNA ASVs were assigned taxonomy based on the Silva version 138 database (Quast et al., 2013), with a minimum bootstrap value of 80/100. ASVs assigned to unknown phyla, mitochondria and chloroplast were removed, and archaeal ASVs were subset into a separate dataset. All phylloplane samples in the archaeal dataset had <10 ASVs and were removed from the analysis. Prior to further analyses, rarefaction curves were generated for all datasets (Fig. S1) which were then rarefied to the minimum sample depth (Table S2) using the *vegan* package version 2.7.1 (Oksanen et al., 2025) . During field sampling, an encounter with aggressive macaques resulted in one sample having an abnormally high relative abundance (> 30%) of Campylobacterota (*Sulfurimonas* and *Arcobacter*) reads compared to all other samples, an indicator of animal contamination, resulting in its removal from further analysis. All visualisations were executed using the *ggplot2* package version 3.5.2 (Wickham, 2016).

### Statistical analyses

Chao1 richness and Pielou’s evenness were calculated for each sample for all gene datasets using the *phyloseq* package version 1.52.0 (McMurdie & Holmes, 2013). The indices were compared between sample types (phylloplane, rhizoplane and sediment) at each location within (early and later succession) and outside the patch using the Kruskal-Wallis test and pairwise Dunn’s tests, performed using the *FSA* package version 0.10.0 (Ogle et al., 2025). A nested linear-mixed effects model with ANOVA testing was fit to the diversity index data to test whether richness or evenness significantly shifted with succession within a specific sample type.

To compare diversity between sediment samples at different positions within the chronosequence and host-associated microbiomes to the above and below ground communities, NMDS ordination and permutation multifactorial analysis of variance (PERMANOVA) tests were performed, based on Bray-Curtis dissimilarity distances. All PERMANOVA analyses used restricted permutations, with sampling patch as a blocking factor, to retain the nested structure of the sampling design. Post-hoc pairwise PERMANOVA testing was performed for the sediment community dataset, for comparisons between succession stages. To assess if diversity within sediments increases along a chronosequence, a linear model was fit to each dataset using both indices and distance of the sample from patch edge.

## RESULTS

Using spatial chronosequence across expanding *Halophila ovalis* patches, we assessed changes in microbial diversity and composition in sediments, rhizoplane and phylloplane communities using taxonomic (16SrRNA gene) and functional (*nifH*, *soxB*, *aprA*, *dsrA*) markers. This was done to uncouple the ecosystem level process of succession from host selection of its microbiome. A chronosequence sampling design is employed to encompass time.

NMDS ordinations of all samples showed strong clustering between sample types (Fig. S2), with distinct community composition between groups across all analysed communities (bacterial, archaeal, *nifH*, *soxB*, *aprA* and *dsrA*) (Table S2).

### Bacterial succession in sediment

Sediments outside the seagrass patch comprise pre-successional microbial communities not yet influenced by a photosynthetic primary coloniser. Between these “pre-succession” sediments and sediments at and within the *H. ovalis* patch edge termed, “early chronosequence sediments”, there are no significant changes in Chao1 richness of ASVs for any of the genes (16S rRNA gene, *nifH, soxB, dsrA,* and *aprA*) (Fig. 2) (Table S3). ASV richness increased significantly between early and later chronosequence sediment for the bacterial 16S rRNA gene and the *nifH* gene (ANOVA, *P* < 0.01) (Fig. 2A & C). Despite clear shifts, this increase is not statistically supported for the archaeal community, nor sulphur cycling genes *soxB*, *aprA* and *dsrA* (Figs. 2B, D-F). Throughout the three successional stages (pre-succession to early to later chronosequence) no significant changes in ASV evenness were detected for any of the genes (Fig. 2) (Table S3).

**Figure 2.**
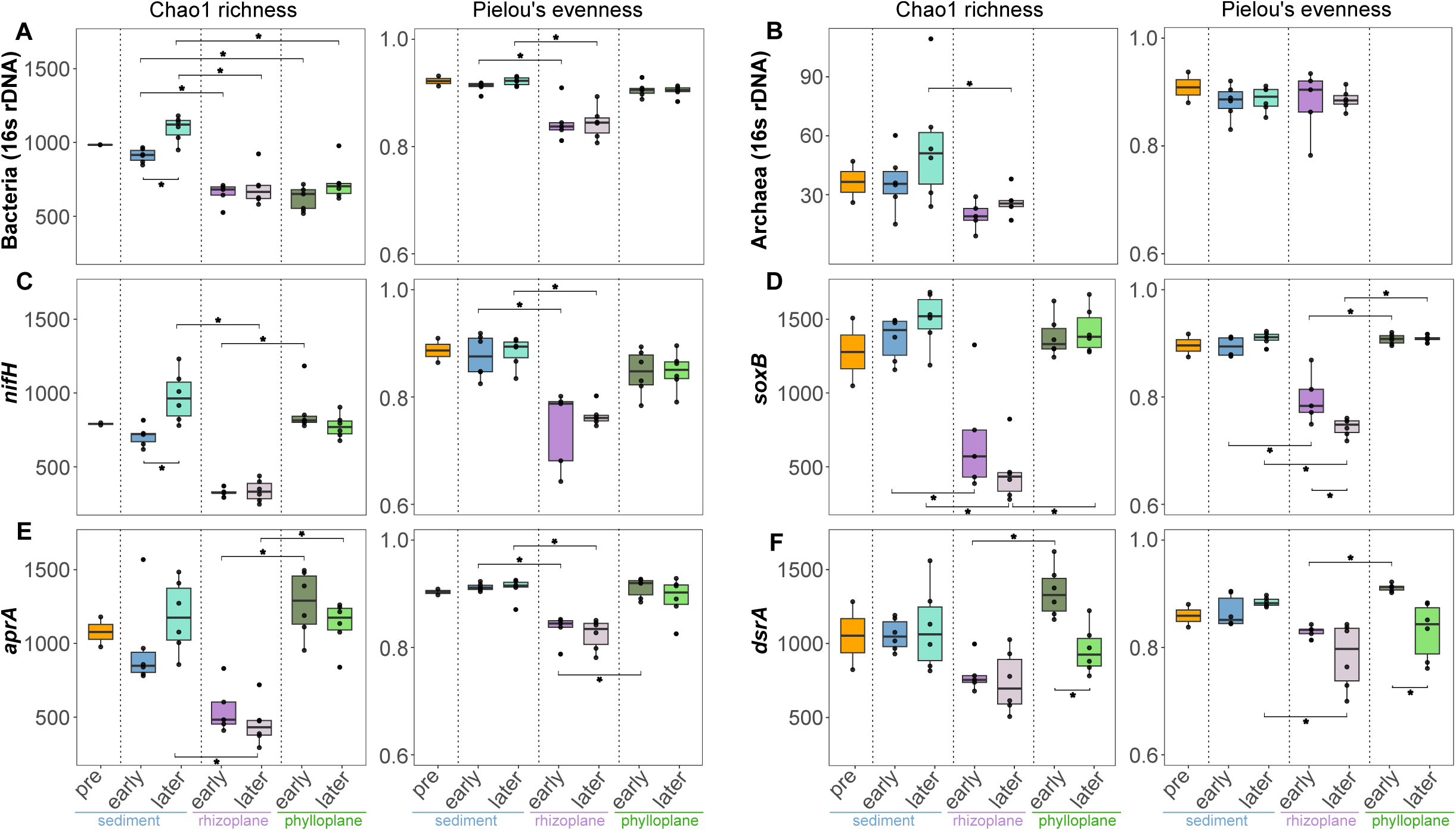
Microbial community richness and evenness. The diversity indices Chao1 richness and Pielou’s evenness are calculated from ASVs of **(A)** bacteria, using the 16s rRNA marker gene, **(B)** archaea, using the 16s rRNA marker gene, and the **(C)** *nifH*, **(D)** *soxB*, **(E)** *aprA* and **(F)** *dsrA* functional genes. Colours of the boxplots correspond to community origin. Statistically significant pair-wise comparisons (Kruskal-Wallis, *P* < 0.05) are represented by asterisks.

Succession is depicted by directional shifts in community composition of habitat forming species, however, succession is frequently inferred from a directional shift in bacterial community composition. This was investigated by comparing community composition for taxonomic and functional gene communities within sediments. They were significantly different between early and later chronosequence sediment communities, for all genes (PERMANOVA, *P* < 0.05), except for the *soxB* (thiosulphate oxidising) functional community (Fig. 3, Table S4).

**Figure 3.**
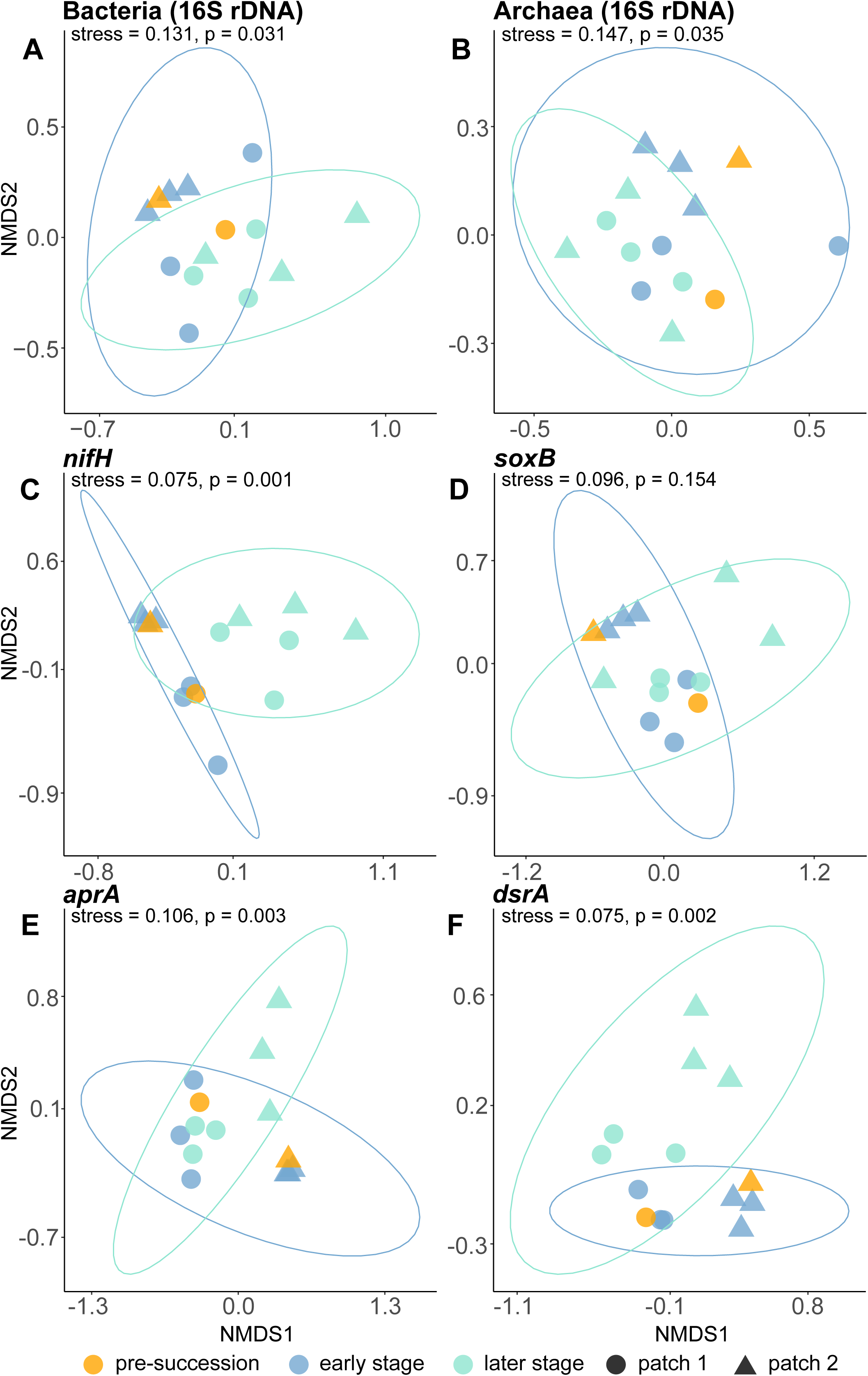
NMDS ordinations of sediment communities between the different stages across the chronosequence. Community composition differences between the three stages of the chronosequence, for (A) bacteria, (B) archaea, (C) *nifH*, (D) *soxB,* (E) *aprA* and (F) *dsrA* communities in the sediment, independent of diversity differences between stages. Colours represent the stages across the chronosequence, while shapes represent the replicate patches sampled.

The association between ASV richness and distance from the seagrass patch edge was investigated, with significant positive correlation detected only in bacteria (linear regression, *P* = 0.001) and *nifH* (*P* = 0.05) communities (Fig. 4) and not for any of the other gene communities (archaeal 16S rRNA gene and S cycling genes: *soxB*, *aprA* and *dsrA* (Fig. S3, Table S5). The correlation for the *nifH* community (Fig. 4B, *R^2^* = 0.26) was weaker than for the bacterial community (Fig. 4A, *R^2^* = 0.63), but still significant. No significant correlation was detected between ASV evenness and distance from the patch edge for any of the genes (Fig. S4, Table S5).

**Figure 4.**
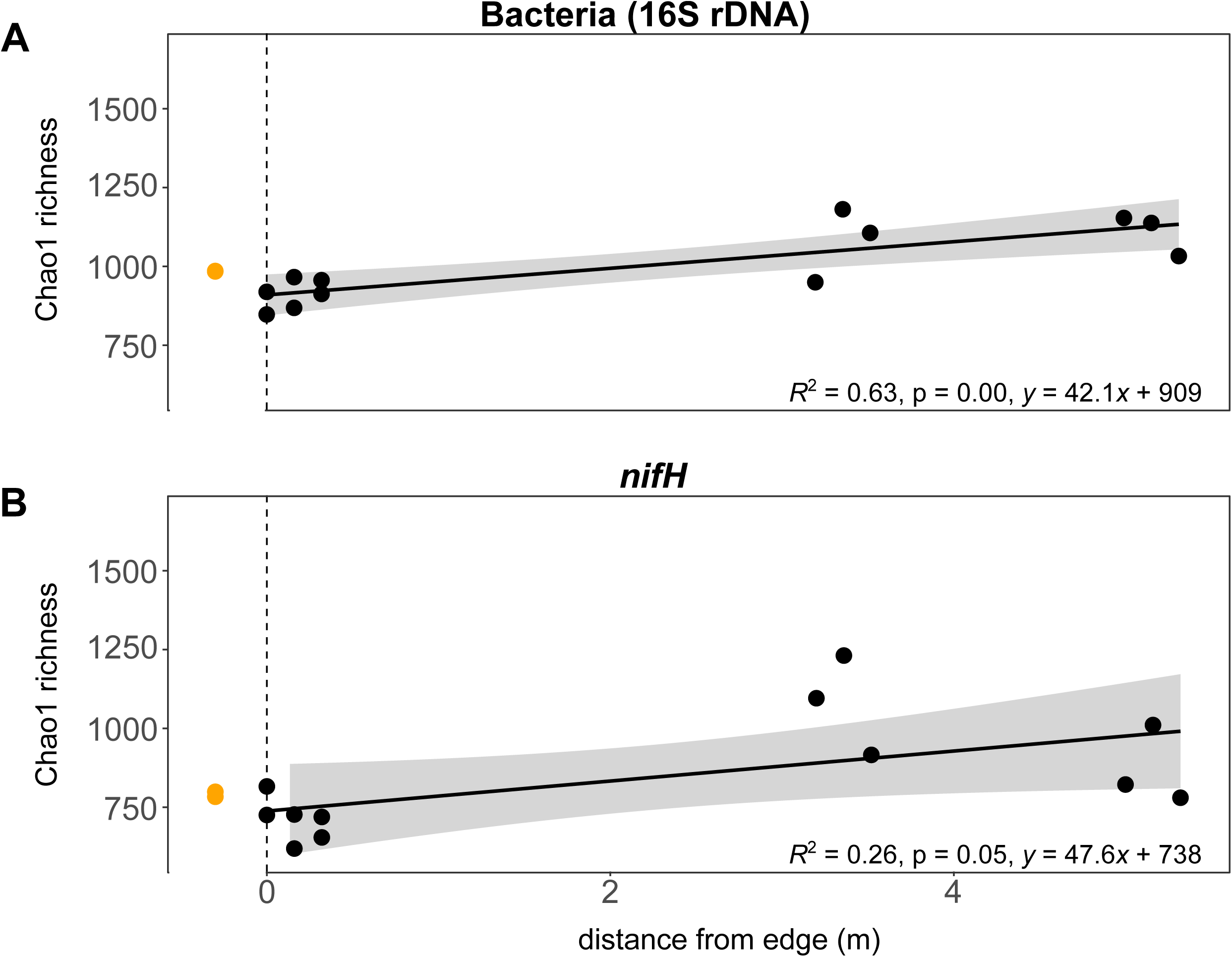
The distance-diversity relationship for the Chao1 richness index, for sediment communities. **(A)** 16S rRNA gene and **(B)** *nifH* analysis of sediment bacterial communities increase in richness with distance from the edge of seagrass growth, as shown with fitted linear regression models. Distance is used here as a substitute for time to study succession, following the chronosequence used in this study. Pre-succession sediment sampled outside the seagrass patch is indicated by the orange points. Grey-shaded areas in the plot indicate the confidence interval (95%) of the best fit line.

### Host selection in seagrass rhizoplane and phylloplane

In early succession, at the edge of the patch and chronosequence, only the bacterial (16S rRNA gene) community richness decreased significantly from sediment to both the organ specific seagrass microbiomes: rhizoplane (root) and phylloplane (leaf) (Kruskal-Wallis, *P* < 0.05, Fig. 2A) (Table S6). *soxB* community also had a significant decrease in ASV richness from sediment to rhizoplane but not to the phylloplane (Fig. 2D) (Table S7). In later succession, the same decreases were detected for bacteria (16S rRNA gene) as in early succession (Kruskal-Wallis, *P* < 0.05, Fig. 2A). Archaea, *nifH*, *soxB* and *aprA* communities also had a significant decrease in ASV richness from sediment to the rhizoplane, but not in the phylloplane (Fig. 2B-E). In terms of evenness, there was a significant decrease from the sediment to rhizoplane communities, for both early and later chronosequence stages (Kruskal-Wallis, *P* < 0.05) for bacteria (16S rRNA gene), *nifH*, *soxB* and *aprA* communities (Fig 2A, C, D, E). For *dsrA*, a lower rhizoplane evenness was observed for the later chronosequence stage only (Fig. 2B & F) (Tables S6-11).

The differences in diversity metrics between seagrass phylloplane and rhizoplane were also calculated. In the early chronosequence stage, ASV richness for *nifH, aprA* and *dsrA* were significantly lower in the rhizoplane than in the phylloplane (Fig. 2C, E & F) while a similar trend was observed in the later chronosequence stage for *soxB* and *aprA* communities (Fig. 2D & E) . Evenness followed a similar trend in the early chronosequence stage, with sulfur cycling genes *soxB*, *aprA* and *dsrA* being significantly lower in the rhizoplane than phylloplane (Kruskal-Wallis, *P* < 0.05, Fig. 2D & E) while this trend was observed only for the *soxB* community in the later chronosequence stage (Fig. 2D) (Tables S6-11).

In contrast to sediments, neither rhizoplane nor phylloplane communities exhibited an increase in ASV richness with succession for any of the genes except *dsrA* phylloplane community where there was a significant decrease over time (ANOVA, *P* < 0.01) (Fig. 2F) (Tables S12-13).

The composition of microbiomes was also investigated for their directional shift in community composition. In contrast to changes in their diversity, community composition had a significant shift from the early to later chronosequence stages, for both the rhizoplane and phylloplane communities, for all genes (PERMANOVA, *P* <0.05) (Figs. S5 & Tables S14-15).

## DISCUSSION

Biofilms are microbial communities, attached to a surface and/or one another, encased in a matrix of biomolecules which creates a habitat of micro-niches. Such communities are distinct from the reversibly attached and motile cells that create the chemically dynamic rhizosphere where diurnal photosynthetic patterns drive the oxic-anoxic gradients of biogeochemical cycling. However, within an ecosystem, organisms are often beyond the direct influence of a host, rather being indirectly influenced by habitat forming species that modify abiotic factors such as sediment quality and structure, water availability, temperature, light and atmospheric chemistry. Such indirect interactions drive ecological phenomena, such as succession. Microbial communities are subject to complex ecological processes and direct species-species interactions. Our study aims to decipher succession and host selection in seagrass meadow formation.

### Community diversity and assemblage trajectories diverge between sediment and host-associated microbiomes

Our results demonstrate that sediment and host-associated microbiomes follow fundamentally different community trajectories during early seagrass patch development, reflecting contrasting assembly processes driven by ecosystem engineering versus microbiomes under host influence. In sediments, increasing bacterial (16S rDNA) and nitrogen-fixing (*nifH*) richness with patch age (Fig. 2) and distance from the patch edge (Fig. 4) is consistent with classical facilitation models of succession, in which habitat-forming organisms progressively modify their environment in ways that permit additional taxa to establish (Connell & Slatyer, 1977; Odum, 1969; Ortiz-Alvarez et al., 2018).

The key findings of bacterial community succession in the seagrass meadow sediments are conceptually synthesized in **Figure 5**. We note that the climax community (Fig. 5A), which would exist within the sediment microbiome of large meadows formed through merging of patches was not sampled in this study due to the complexity of other ecological processes that drive dynamic changes rather than increasing diversity (illustrated as a pale green wedge, Fig. 5A).

**Figure 5.**
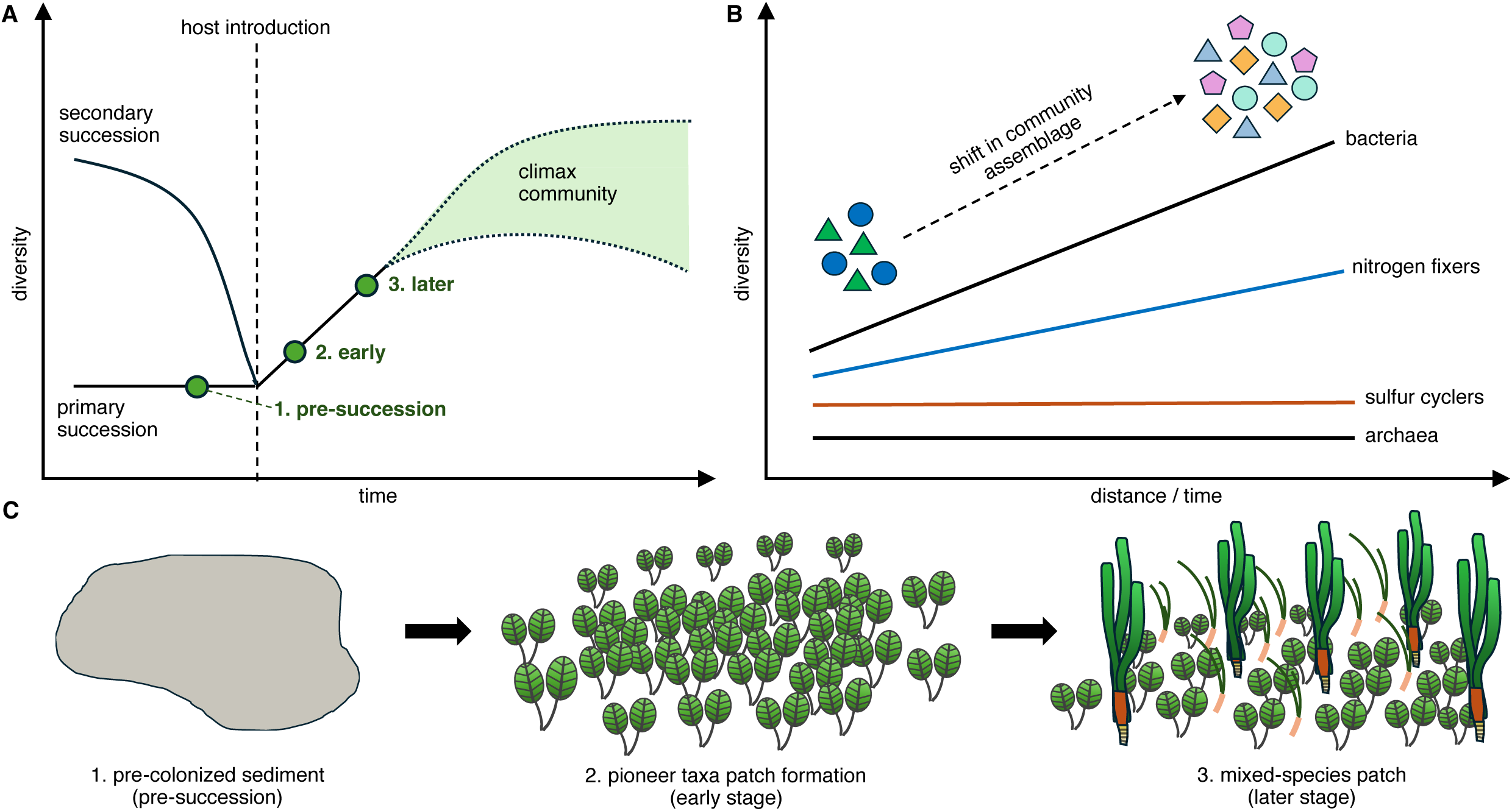
Stages and diversity measurements of a seagrass meadow ecological succession model. A chronosequence from expanding seagrass patches was sampled along. **(A)** three stages of the chronosequence (pre-succession, early and later) that were sampled from a theoretical succession model that demonstrates the increase in microbial diversity upon introduction of a habitat-forming host. Pre-climax communities were sampled as climax communities undergo varying ecological dynamics that produce a diversity range (pale green). **(B)** Significant increase in alpha diversity for bacteria and nitrogen fixing communities along the chronosequence, differentially effected to sulphur cycling communities. Changes in beta diversity was detected for all communities. **(C)** Visual representation of the three stages of seagrass patch expansion that was sampled.

Mechanistically, this diversification is driven by the accumulation of plant-derived organic matter, including particulate detritus from senescing leaves and roots, dissolved organic carbon released via root exudation, and labile carbohydrates such as sucrose that accumulate at exceptionally high concentrations in seagrass sediments (Duarte et al., 2010; Hennion et al., 2019; Sogin et al., 2022). These carbon pools fuel heterotrophic metabolism and increase substrate heterogeneity, thereby supporting a higher diversity of metabolically and behaviourally diverse microbes as the patches mature (Kuzyakov & Razavi, 2019).

Seagrass expansion also generates steep and persistent redox gradients within sediments. This begins when a growing root first penetrates marine sediments and there is a concentrated and localised release of oxygen, immediately lowering the concentration of toxic reduced sulphur cycle intermediates (Martin et al., 2018) and stimulating microbes behaviourally responsive (Seymour et al., 2017) and metabolically capable of aerobic metabolism. Oxygen released from roots during photosynthesis produce localised oxic microzones embedded within anoxic sediments (Fig. 1D), while diel cycles of oxygen release and consumption create temporally dynamic redox conditions (Brodersen et al., 2015; Hasler-Sheetal & Holmer, 2015; Sand-Jensen et al., 2005). These gradients allow the coexistence of aerobic heterotrophs, facultative anaerobes, sulfate reducers, sulfur oxidisers, and nitrogen fixers across fine spatial scales, promoting compositional turnover as successional time increases (Ashkaan K. Fahimipour et al., 2017; Jiang et al., 2016). Importantly, sustained rhizodeposition, which is the continuous release of root-derived carbon compounds, oxygen, and metabolites into sediments over the lifetime of a seagrass patch drives sediment modification (Kuzyakov & Razavi, 2019; Sun et al., 2025). Thus from our chronosequence, we infer that as biomass of *H. ovalis* increases, these inputs become increasingly spatially extensive and temporally stable, resulting in cumulative exposure of sediment microbial communities to root-mediated processes (Tomlinson, 1974; Walker et al., 2010). The observed increase in bacterial and *nifH* richness with distance from the patch edge (Fig. 4) therefore reflects time-integrated ecosystem engineering rather than simple spatial heterogeneity.

In contrast, host-associated microbiomes (i.e. surface attached biofilms) on the rhizoplane and phylloplane did not exhibit increasing richness or evenness with succession (Fig. 2), despite significant shifts in community composition (Figs S5 and S6). This pattern indicates that host-associated communities are not assembled through facilitation-driven diversification but are instead shaped by strong host-mediated environmental filtering, consistent with holobiont theory (Simon et al., 2019; Tarquinio et al., 2019). The seagrass host constrains microbial retainment through oxygen regulation, carbon allocation, antimicrobial compounds, chemotactic cues and repellents, and immune-like recognition processes (Sogin et al., 2022; Ugarelli et al., 2017; Vogel et al., 2020), resulting in early stabilisation of diversity followed by compositional replacement rather than expansion. We suggest that the stronger filtering observed in the rhizoplane relative to the phylloplane (Fig. 2) further reflects functional asymmetry within the holobiont. Below-ground tissues experience persistent anoxia, sulfide exposure, and strong redox oscillations, necessitating tight regulation of microbial partners involved in detoxification and nutrient exchange (Hasler-Sheetal & Holmer, 2015). As a result, rhizoplane communities converge toward low-diversity, host-selected assemblages even as sediment communities continue to diversify.

### Nitrogen fixation diversity scales with taxonomic diversity across succession, but sulfur cycling diversity does not

Across the five genes examined, functional diversity of sulfur cyclers (*soxB*, *aprA*, *dsrA*) did not scale linearly with taxonomic diversity (16S rDNA), demonstrating a decoupling between organismal richness and biogeochemical potential during succession. In sediments, increases in bacterial and *nifH* ASV richness were not accompanied by parallel increases in sulfur-cycling ASV richness (Fig. 2), indicating that expansion of taxonomic diversity does not necessarily translate into functional diversification (Louca et al., 2018). This decoupling reflects fundamental constraints on microbial metabolism. Sulfate reduction and sulfur oxidation are governed primarily by thermodynamics and redox chemistry, limiting the range of viable metabolic pathways despite taxonomic turnover (Jørgensen, 1982; Muyzer & Stams, 2008). Consequently, sulfur-cycling communities often exhibit high functional redundancy, with succession proceeding through replacement among ecologically equivalent taxa rather than functional expansion (Ashkaan K. Fahimipour et al., 2017; Hasler-Sheetal & Holmer, 2015). The stability of *dsrA*, *aprA*, and *soxB* ASV richness across successional stages in our study is consistent with this model (Fig. S3).

The *dsrA* and *aprA* genes occur in both sulfate-reducing and sulfur-oxidizing microbes, where they function in distinct reductive and oxidative pathways, leading them to be described as reversible, whereas *soxB* functions unidirectionally in sulfur oxidation. All three sulfur-cycling genes generally exhibit reduced diversity along the chronosequence, indicating strong host selection. In contrast, *nifH* diversity in rhizoplane and phylloplane remains stable, suggesting that nitrogen-fixers are maintained rather than excluded. The similar reduction in diversity across sulfur genes, regardless of whether they participate in oxidative or reductive pathways, indicates that selection is not structured by reaction bidirectionality. Instead, the host increasingly selects, along the chronosequence, a subset of sulfur-cycling taxa involved in detoxification and redox regulation, while nitrogen fixation remains as a stable function.

Seagrasses actively regulate microbial metabolism through oxygen production and transport from leaf to root, where it exerts stronger selection within a low (root) versus high (leaf) redox niche. In addition, seagrasses exert selective carbon provisioning, and host-derived chemical cues, thereby favouring microbes that perform essential services while excluding functionally incompatible taxa (Tarquinio et al., 2019; Ugarelli et al., 2017). This results in early establishment of a functionally coherent microbiome that persists through successional time. Nitrogen fixation illustrates this selective constraint particularly clearly. While *nifH* diversity increased with sediment succession, likely driven by expanding carbon availability and microscale anoxia, *nifH* richness remained unchanged between early and later stages in both rhizoplane and phylloplane. This suggests that seagrass roots selectively recruit and regulate nitrogen-fixing partners rather than passively tracking sedimentary diversification (Crump et al., 2018; Mohr et al., 2021), while *nifH* richness in the phylloplane is unchanging. This lack of dynamism suggests that nitrogen fixers once inoculated as the seagrass grows through sediment, remains stable despite the aerobic intertidal dynamic of ocean and air submersion. Given the high energetic cost and oxygen sensitivity of nitrogen fixation, such regulation likely reflects a balance between nitrogen acquisition and metabolic risk to the host.

### Succession differentially influences taxonomic and biogeochemical community structure

Comparisons between taxonomic and functional assemblages (Fig. 3, S5 and S6) reveal that succession exerts a stronger influence on community composition than on functional diversity. Across sediments, rhizoplane, and phylloplane, significant compositional shifts occurred between early and later stages for all gene communities examined, yet these shifts were rarely accompanied by increases in functional richness. Together, these results show that succession mainly involves changes in which taxa perform existing functions, rather than the emergence of new microbial functions (Allison & Martiny, 2008; Louca et al., 2018). This dynamic is especially evident in sulfur-cycling communities, where stable functional richness alongside compositional turnover suggests that key detoxification pathways are buffered against environmental change. Such buffering is critical in seagrass ecosystems, where sulfide accumulation poses a persistent threat to plant survival (Hasler-Sheetal & Holmer, 2015; Jiang et al., 2016). Functional redundancy among sulfur-cycling microbes ensures continuity of sulfide oxidation and reduction even as community composition reorganises across successional stages.

Within the rhizoplane and phylloplane biofilm, this functional stability is consistent with predictions of holobiont theory, which posit that hosts stabilise essential microbial functions while allowing flexibility in taxonomic membership to enhance resilience (Rosenberg & Zilber-Rosenberg, 2018; Simon et al., 2019). Our results support this framework, showing that while the assemblage of host-associated microbial taxa shift with patch age, the functional roles they perform remain conserved. Indeed, while changes in taxonomic and functional assemblages do accompany succession, changes in assemblage can result from many ecological phenomena, such as host selection, and therefore change in assemblage cannot be indicative of succession.

Together, these findings reconcile classical succession theory with host-microbe dynamics by demonstrating that microbial succession in seagrass ecosystems is compartment-specific. Sediments diversify through cumulative ecosystem engineering that increases physicochemical heterogeneity, whereas host-associated microbiomes are assembled through strong filtering that stabilises function while permitting taxonomic turnover. Succession determines which taxa are available, but host selection of its microbiome’s assembly determines which functions persist.

## CONCLUSION

By integrating the 16S rRNA taxonomic marker with multiple functional genes spanning nitrogen and sulfur cycling, this study demonstrates that early seagrass succession is characterised by spatially partitioned assembly regimes along a chronosequence. The surrounding sediment acts as a successional arena where ecosystem engineering drives increasing taxonomic diversity, while the seagrass microbiome functions as a selected community at the host surface’s interface, maintaining functional processes beneficial to the host. This duality provides a mechanistic framework for understanding how seagrass meadows maintain biogeochemical stability and drive functional dynamics during expansion and recovery in increasingly disturbed coastal environments.

## ACKNOWLEDGEMENTS

We thank Su Xuan for assistance with field sampling and support from the National Parks Board (NP_RP23-095). This study, CWHS and SS are supported by the Marine Climate Change Science Programme (MCCS Award NRF-MCCS21-1-4-0002) under the National Research Foundation, Singapore. The Singapore Centre for Environmental Life Sciences Engineering (SCELSE) is funded by the Ministry of Education, Singapore, the National Research Foundation of Singapore, Nanyang Technological University Singapore, and the National University of Singapore.

## AUTHOR CONTRIBUTIONS

RJC contributed to the study’s conception and design. PM contributed to the experimental design and acquired the samples. PM and KZC processed the samples, generating data. PM, CWHS, SS and RJC analysed and interpreted the data. CWHS and PM drafted the manuscript. All authors have revised and approved the final manuscript.

## DATA AVAILABILITY

Raw sequencing data have been deposited at the NCBI Sequence Read Archive (SRA) under the accession number PRJNA1398339.

## References cited

1. Aasfar, A., Bargaz, A., Yaakoubi, K., Hilali, A., Bennis, I., Zeroual, Y., & Meftah Kadmiri, I. (2021). Nitrogen fixing *Azotobacter* species as potential soil biological enhancers for crop nutrition and yield stability. Frontiers in Microbiology, 12, 628379. 10.3389/fmicb.2021.628379

2. Allison, S. D., & Martiny, J. B. H. (2008). Resistance, resilience, and redundancy in microbial communities. Proceedings of the National Academy of Sciences, 105(supplement_1), 11512-11519. doi:10.1073/pnas.0801925105

3. Aoki, M., Kakiuchi, R., Yamaguchi, T., Takai, K., Inagaki, F., & Imachi, H. (2015). Phylogenetic diversity of *aprA* genes in subseafloor sediments on the Northwestern Pacific margin off Japan. Microbes Environ, 30(3), 276–280. 10.1264/jsme2.ME15023

4. Banister, R. B., Schwarz, M. T., Fine, M., Ritchie, K. B., & Muller, E. M. (2022). Instability and stasis among the microbiome of seagrass leaves, roots and rhizomes, and nearby sediments within a natural pH gradient. Microb Ecol, 84(3), 703–716. 10.1007/s00248-021-01867-9

5. Barea, J.-M., Pozo, M. J., Azcón, R., & Azcón-Aguilar, C. (2005). Microbial co-operation in the rhizosphere. Journal of Experimental Botany, 56(417), 1761–1778. 10.1093/jxb/eri197

6. Brodersen, K. E., Mosshammer, M., Bittner, M. J., Hallstrøm, S., Santner, J., Riemann, L., & Kühl, M. (2024). Seagrass-mediated rhizosphere redox gradients are linked with ammonium accumulation driven by diazotrophs. Microbiol Spectr, 12(4), e0333523. 10.1128/spectrum.03335-23

7. Brodersen, K. E., Nielsen, D. A., Ralph, P. J., & Kühl, M. (2015). Oxic microshield and local pH enhancement protects Zostera muelleri from sediment derived hydrogen sulphide. New Phytologist, 205(3), 1264–1276. 10.1111/nph.13124

8. Brodersen, K. E., Siboni, N., Nielsen, D. A., Pernice, M., Ralph, P. J., Seymour, J., & Kühl, M. (2018). Seagrass rhizosphere microenvironment alters plant-associated microbial community composition. Environ Microbiol, 20(8), 2854–2864. 10.1111/1462-2920.14245

9. Callahan, B. J., McMurdie, P. J., Rosen, M. J., Han, A. W., Johnson, A. J., & Holmes, S. P. (2016). DADA2: High-resolution sample inference from Illumina amplicon data. Nat Methods, 13(7), 581–583. 10.1038/nmeth.3869

10. Caporaso, J. G., Lauber, C. L., Walters, W. A., Berg-Lyons, D., Lozupone, C. A., Turnbaugh, P. J., Fierer, N., & Knight, R. (2011). Global patterns of 16S rRNA diversity at a depth of millions of sequences per sample. Proc Natl Acad Sci U S A, 108 Suppl 1(Suppl 1), 4516–4522. 10.1073/pnas.1000080107

11. Connell, J. H., & Slatyer, R. O. (1977). Mechanisms of succession in natural communities and their role in community stability and organization. The American naturalist, 111(982), 1119–1144. 10.1086/283241

12. Crump, B. C., Wojahn, J. M., Tomas, F., & Mueller, R. S. (2018). Metatranscriptomics and Amplicon Sequencing Reveal Mutualisms in Seagrass Microbiomes. Frontiers in Microbiology, 9, 388. 10.3389/fmicb.2018.00388

13. Duarte, C. M., Marbà, N., Gacia, E., Fourqurean, J. W., Beggins, J., Barrón, C., & Apostolaki, E. T. (2010). Seagrass community metabolism: Assessing the carbon sink capacity of seagrass meadows. Global Biogeochemical Cycles, 24(4). 10.1029/2010GB003793

14. Fahimipour, A. K., Kardish, M. R., Lang, J. M., Green, J. L., Eisen, J. A., & Stachowicz, J. J. (2017). Global-Scale Structure of the Eelgrass Microbiome. Appl Environ Microbiol, 83(12). 10.1128/aem.03391-16

15. Fahimipour, A. K., Kardish, M. R., Lang, J. M., Green, J. L., Eisen, J. A., & Stachowicz, J. J. (2017). Global-scale structure of the Eelgrass microbiome. Applied and Environmental Microbiology, 83(12), e03391–03316. doi:10.1128/AEM.03391-16

16. Fortes, M. D., Ooi, J. L. S., Tan, Y. M., Prathep, A., Bujang, J. S., & Yaakub, S. M. (2018). Seagrass in Southeast Asia: a review of status and knowledge gaps, and a road map for conservation. Botanica Marina, 61(3), 269–288. doi:10.1515/bot-2018-0008

17. Friess, D. A., Gatt, Y. M., Fung, T. K., Alemu, J. B., Bhatia, N., Case, R., Chua, S. C., Huang, D., Kwan, V., Lim, K. E., Nathan, Y., Ow, Y. X., Saavedra-Hortua, D., Sloey, T. M., Yando, E. S., Ibrahim, H., Koh, L. P., Puah, J. Y., Teo, S. L.-M., Tun, K., Wong, L. W., & Yaakub, S. M. (2023). Blue carbon science, management and policy across a tropical urban landscape. Landscape and Urban Planning, 230, 104610. 10.1016/j.landurbplan.2022.104610

18. Gao, H., Wang, C., Chen, J., Wang, P., Zhang, J., Zhang, B., Wang, R., & Wu, C. (2022). Enhancement effects of decabromodiphenyl ether on microbial sulfate reduction in eutrophic lake sediments: A study on sulfate-reducing bacteria using *dsrA* and *dsrB* amplicon sequencing. Science of The Total Environment, 843, 157073. 10.1016/j.scitotenv.2022.157073

19. García-Martínez, M., Kuo, J., Kilminster, K., Walker, D., Rosselló-Mora, R., & Duarte, C. M. (2005). Microbial colonization in the seagrass Posidonia spp. roots. Marine Biology Research, 1(6), 388–395. 10.1080/17451000500443419

20. Gauri, S. S., Mandal, S. M., & Pati, B. R. (2012). Impact of Azotobacter exopolysaccharides on sustainable agriculture. Applied Microbiology and Biotechnology, 95(2), 331–338. 10.1007/s00253-012-4159-0

21. Hasler-Sheetal, H., & Holmer, M. (2015). Sulfide intrusion and detoxification in the seagrass Zostera marina. PLoS One, 10(6), e0129136. 10.1371/journal.pone.0129136

22. Hennion, N., Durand, M., Vriet, C., Doidy, J., Maurousset, L., Lemoine, R., & Pourtau, N. (2019). Sugars en route to the roots. Transport, metabolism and storage within plant roots and towards microorganisms of the rhizosphere. Physiologia Plantarum, 165(1), 44–57. https://doi-org.remotexs.ntu.edu.sg/10.1111/ppl.12751

23. Hou, J., Sievert, S. M., Wang, Y., Seewald, J. S., Natarajan, V. P., Wang, F., & Xiao, X. (2020). Microbial succession during the transition from active to inactive stages of deep-sea hydrothermal vent sulfide chimneys. Microbiome, 8(1), 102. 10.1186/s40168-020-00851-8

24. Iqbal, M. M., Nishimura, M., Haider, M. N., & Yoshizawa, S. (2022). Microbial communities on eelgrass (*Zostera marina*) thriving in Tokyo Bay and the possible source of leaf-attached microbes. Front Microbiol, 13, 1102013. 10.3389/fmicb.2022.1102013

25. Jensen, S. I., Kühl, M., & Priemé, A. (2007). Different bacterial communities associated with the roots and bulk sediment of the seagrass Zostera marina. FEMS Microbiology Ecology, 62(1), 108–117. 10.1111/j.1574-6941.2007.00373.x

26. Jiang, J., Chan, A., Ali, S., Saha, A., Haushalter, K. J., Lam, W. L., Glasheen, M., Parker, J., Brenner, M., Mahon, S. B., Patel, H. H., Ambasudhan, R., Lipton, S. A., Pilz, R. B., & Boss, G. R. (2016). Hydrogen sulfide - mechanisms of toxicity and development of an antidote. Scientific Reports, 6, 20831. 10.1038/srep20831

27. Jørgensen, B. B. (1982). Mineralization of organic matter in the sea bed—the role of sulphate reduction. Nature, 296(5858), 643–645. 10.1038/296643a0

28. Kuzyakov, Y., & Razavi, B. S. (2019). Rhizosphere size and shape: Temporal dynamics and spatial stationarity. Soil Biology and Biochemistry, 135, 343–360. 10.1016/j.soilbio.2019.05.011

29. Kwan, V., Shantti, P., Lum, E. Y. Y., Ow, Y. X., & Huang, D. (2023). Diversity and phylogeny of seagrasses in Singapore. Aquatic Botany, 187. 10.1016/j.aquabot.2023.103648

30. Lamb, J. B., van de Water, J. A. J. M., Bourne, D. G., Altier, C., Hein, M. Y., Fiorenza, E. A., Abu, N., Jompa, J., & Harvell, C. D. (2017). Seagrass ecosystems reduce exposure to bacterial pathogens of humans, fishes, and invertebrates. Science, 355(6326), 731–733. doi:10.1126/science.aal1956

31. Longford, S. R., Campbell, A. H., Nielsen, S., Case, R. J., Kjelleberg, S., & Steinberg, P. D. (2019). Interactions within the microbiome alter microbial interactions with host chemical defences and affect disease in a marine holobiont. Scientific Reports, 9(1), 1363. 10.1038/s41598-018-37062-z

32. Louca, S., Polz, M. F., Mazel, F., Albright, M. B. N., Huber, J. A., O’Connor, M. I., Ackermann, M., Hahn, A. S., Srivastava, D. S., Crowe, S. A., Doebeli, M., & Parfrey, L. W. (2018). Function and functional redundancy in microbial systems. Nature Ecology & Evolution, 2(6), 936–943. 10.1038/s41559-018-0519-1

33. Majzoub, M. E., Beyersmann, P. G., Simon, M., Thomas, T., Brinkhoff, T., & Egan, S. (2019). *Phaeobacter inhibens* controls bacterial community assembly on a marine diatom. FEMS Microbiol Ecol, 95(6). 10.1093/femsec/fiz060

34. Martin, B. C., Bougoure, J., Ryan, M. H., Bennett, W. W., Colmer, T. D., Joyce, N. K., Olsen, Y. S., & Kendrick, G. A. (2018). Oxygen loss from seagrass roots coincides with colonisation of sulphide-oxidising cable bacteria and reduces sulphide stress. The ISME Journal, 13(3), 707–719. 10.1038/s41396-018-0308-5

35. Martin, B. C., Bougoure, J., Ryan, M. H., Bennett, W. W., Colmer, T. D., Joyce, N. K., Olsen, Y. S., & Kendrick, G. A. (2019). Oxygen loss from seagrass roots coincides with colonisation of sulphide-oxidising cable bacteria and reduces sulphide stress. ISME J, 13(3), 707–719. 10.1038/s41396-018-0308-5

36. Martin, M. (2011). Cutadapt removes adapter sequences from high throughput sequencing reads. EMBnet Journal, 17, 10–12. 10.14806/ej.17.1.200

37. Mazarrasa, I., Marbà, N., Lovelock, C. E., Serrano, O., Lavery, P. S., Fourqurean, J. W., Kennedy, H., Mateo, M. A., Krause-Jensen, D., Steven, A. D. L., & Duarte, C. M. (2015). Seagrass meadows as a globally significant carbonate reservoir. Biogeosciences, 12(16), 4993–5003. 10.5194/bg-12-4993-2015

38. McKenzie, L. J., Yaakub, S. M., Tan, R., Seymour, J., & Yoshida, R. L. (2016). Seagrass habitats of Singapore: Environmental drivers and key processes. Raffles Bulletin of Zoology, 60–77.

39. McMurdie, P. J., & Holmes, S. H. (2013). phyloseq: An R package for reproducible interactive analysis and graphics of microbiome census data. PLoS One, 8, e61217. http://dx.plos.org/10.1371/journal.pone.0061217

40. Mohr, W., Lehnen, N., Ahmerkamp, S., Marchant, H. K., Graf, J. S., Tschitschko, B., Yilmaz, P., Littmann, S., Gruber-Vodicka, H., Leisch, N., Weber, M., Lott, C., Schubert, C. J., Milucka, J., & Kuypers, M. M. M. (2021). Terrestrial-type nitrogen-fixing symbiosis between seagrass and a marine bacterium. Nature, 600(7887), 105–109. 10.1038/s41586-021-04063-4

41. Mönnich, J., Tebben, J., Bergemann, J., Case, R., Wohlrab, S., & Harder, T. (2020). Niche-based assembly of bacterial consortia on the diatom *Thalassiosira rotula* is stable and reproducible. The ISME Journal, 14(6), 1614–1625. 10.1038/s41396-020-0631-5

42. Mushegian, A. A., Peterson, C. N., Baker, C. C., & Pringle, A. (2011). Bacterial diversity across individual lichens. Applied Environmental Microbiology, 77(12), 4249–4252. 10.1128/AEM.02850-10

43. Muyzer, G., & Stams, A. J. M. (2008). The ecology and biotechnology of sulphate-reducing bacteria. Nature Reviews Microbiology, 6(6), 441–454. 10.1038/nrmicro1892

44. Odum, E. P. (1969). The strategy of ecosystem development. Science, 164(3877), 262–270. doi:10.1126/science.164.3877.262

45. Ogle, D. H., Doll, J. C., Wheeler, A. P., & Dinno, A. (2025). FSA: Simple Fisheries Stock Assessment Methods. 10.32614/CRAN.package.FSA

46. Oksanen, J., Simpson, G., Blanchet, F., Kindt, R., Legendre, P., Minchin, P., O’Hara, R., Solymos, P., Stevens, M., Szoecs, E., Wagner, H., Barbour, M., Bedward, M., Bolker, B., Borcard, D., Borman, T., Carvalho, G., Chirico, M., De Caceres, M., Durand, S., Evangelista, H., FitzJohn, R., Friendly, M., Furneaux, B., Hannigan, G., Hill, M., Lahti, L., Martino, C., McGlinn, D., Ouellette, M., Ribeiro Cunha, E., Smith, T., Stier, A., Ter Braak, C., & Weedon, J. (2025). vegan: Community Ecology Package. 10.32614/CRAN.package.vegan

47. Ortiz-Alvarez, R., Fierer, N., de Los Rios, A., Casamayor, E. O., & Barberan, A. (2018). Consistent changes in the taxonomic structure and functional attributes of bacterial communities during primary succession. The ISME Journal, 12(7), 1658–1667. 10.1038/s41396-018-0076-2

48. Petri, R., Podgorsek, L., & Imhoff, J. F. (2001). Phylogeny and distribution of the *soxB* gene among thiosulfate-oxidizing bacteria. FEMS Microbiol Lett, 197(2), 171–178. 10.1111/j.1574-6968.2001.tb10600.x

49. Poly, F., Monrozier, L. J., & Bally, R. (2001). Improvement in the RFLP procedure for studying the diversity of *nifH* genes in communities of nitrogen fixers in soil. Res Microbiol, 152(1), 95–103. 10.1016/s0923-2508(00)01172-4

50. Polz, M. F., & Cavanaugh, C. M. (1998). Bias in template-to-product ratios in multitemplate PCR. Appl Environ Microbiol, 64(10), 3724–3730. 10.1128/aem.64.10.3724-3730.1998

51. Quast, C., Pruesse, E., Yilmaz, P., Gerken, J., Schweer, T., Yarza, P., Peplies, J., & Glockner, F. O. (2013). The SILVA ribosomal RNA gene database project: improved data processing and web-based tools. Nucleic Acids Res, 41(Database issue), D590–596. 10.1093/nar/gks1219

52. R Core Team. (2025). R: A Language and Environment for Statistical Computing. https://www.R-project.org/

53. Rabbani, G., Yan, B. C., Lee, N. L. Y., Ooi, J. L. S., Lee, J. N., Huang, D., & Wainwright, B. J. (2021). Spatial and structural factors shape seagrass-associated bacterial communities in Singapore and peninsular Malaysia. Frontiers in Marine Science, 8. 10.3389/fmars.2021.659180

54. Rosenberg, E., & Zilber-Rosenberg, I. (2018). The hologenome concept of evolution after 10 years. Microbiome, 6(1), 78. 10.1186/s40168-018-0457-9

55. Sand-Jensen, K., Pedersen, O., Binzer, T., & Borum, J. (2005). Contrasting oxygen dynamics in the freshwater isoetid Lobelia dortmanna and the marine seagrass Zostera marina. Annals of Botany, 96(4), 613–623. 10.1093/aob/mci214

56. Seymour, J. R., Amin, S. A., Raina, J.-B., & Stocker, R. (2017). Zooming in on the phycosphere: the ecological interface for phytoplankton–bacteria relationships. Nature Microbiology, 2(7), 17065. 10.1038/nmicrobiol.2017.65

57. Simon, J.-C., Marchesi, J. R., Mougel, C., & Selosse, M.-A. (2019). Host-microbiota interactions: from holobiont theory to analysis. Microbiome, 7(1), 5. 10.1186/s40168-019-0619-4

58. Sogin, E. M., Michellod, D., Gruber-Vodicka, H. R., Bourceau, P., Geier, B., Meier, D. V., Seidel, M., Ahmerkamp, S., Schorn, S., D’Angelo, G., Procaccini, G., Dubilier, N., & Liebeke, M. (2022). Sugars dominate the seagrass rhizosphere. Nature Ecology & Evolution, 6(7), 866–877. 10.1038/s41559-022-01740-z

59. Stephens, B. M., Durkin, C. A., Sharpe, G., Nguyen, T. T. H., Albers, J., Estapa, M. L., Steinberg, D. K., Levine, N. M., Gifford, S. M., Carlson, C. A., Boyd, P. W., & Santoro, A. E. (2024). Direct observations of microbial community succession on sinking marine particles. The ISME Journal, 18(1). 10.1093/ismejo/wrad010

60. Sun, J., Zhao, X., Liu, J.-J., Zhang, Z., Yan, W.-J., & Zhang, P.-D. (2025). Functional specialization of the eelgrass rhizosphere microbiome: Root exudate-mediated assembly and implications for degraded meadow rehabilitation. Marine Environmental Research, 212, 107577. 10.1016/j.marenvres.2025.107577

61. Tarquinio, F., Hyndes, G. A., Laverock, B., Koenders, A., & Säwström, C. (2019). The seagrass holobiont: understanding seagrass-bacteria interactions and their role in seagrass ecosystem functioning. FEMS Microbiology Letters, 366(6). 10.1093/femsle/fnz057

62. Thompson, J. R., Marcelino, L. A., & Polz, M. F. (2002). Heteroduplexes in mixed-template amplifications: formation, consequence and elimination by ’reconditioning PCR’. Nucleic Acids Res, 30(9), 2083–2088. 10.1093/nar/30.9.2083

63. Thomson, J. A., Burkholder, D. A., Heithaus, M. R., Fourqurean, J. W., Fraser, M. W., Statton, J., & Kendrick, G. A. (2015). Extreme temperatures, foundation species, and abrupt ecosystem change: an example from an iconic seagrass ecosystem. Global Change Biology, 21(4), 1463–1474. 10.1111/gcb.12694

64. Tomlinson, P. B. (1974). Vegetative morphology and meristem dependence — The foundation of productivity in seagrasses. Aquaculture, 4, 107–130. 10.1016/0044-8486(74)90027-1

65. Ugarelli, K., Chakrabarti, S., Laas, P., & Stingl, U. (2017). The Seagrass Holobiont and Its Microbiome. Microorganisms, 5(4), 81. https://www.mdpi.com/2076-2607/5/4/81

66. Vogel, M. A., Mason, O. U., & Miller, T. E. (2020). Host and environmental determinants of microbial community structure in the marine phyllosphere. PLoS One, 15(7), e0235441. 10.1371/journal.pone.0235441

67. Walker, L. R., Wardle, D. A., Bardgett, R. D., & Clarkson, B. D. (2010). The use of chronosequences in studies of ecological succession and soil development. Journal of Ecology, 98(4), 725–736. 10.1111/j.1365-2745.2010.01664.x

68. Ward, M., Kindinger, T. L., Hirsh, H. K., Hill, T. M., Jellison, B. M., Lummis, S., Rivest, E. B., Waldbusser, G. G., Gaylord, B., & Kroeker, K. J. (2022). Reviews and syntheses: Spatial and temporal patterns in seagrass metabolic fluxes. Biogeosciences, 19(3), 689–699. 10.5194/bg-19-689-2022

69. Wickham, H. (2016). ggplot2: Elegant Graphics for Data Analysis. Springer-Verlag New York. https://ggplot2.tidyverse.org

70. Yaakub, S. M., McKenzie, L. J., Erftemeijer, P. L. A., Bouma, T., & Todd, P. A. (2014). Courage under fire: Seagrass persistence adjacent to a highly urbanised city–state. Marine Pollution Bulletin, 83(2), 417–424. 10.1016/j.marpolbul.2014.01.012

71. Yan, B. C., Rabbani, G., Lee, N. L. Y., Ooi, J. L. S., Lee, J. N., Huang, D., & Wainwright, B. J. (2021). The microbiome of the seagrass *Halophila ovalis*: community structuring from plant parts to regional scales. Aquatic Microbial Ecology, 87, 139–150. 10.3354/ame01976

