## Supplementary Materials for "Uncoupling of seagrass host selection and succession for microbial guilds in meadow chronosequence"

**SUPPLEMENTARY TABLES**

**Table S1. PCR conditions for each amplicon analysed in this study.** All amplified fragment lengths exclude primers. A total reaction volume of 25 µl, and 25 cycles were used for the 16S rRNA marker gene. A total reaction volume of 50 µl, and 30 cycles were used for the functional genes. PCR steps repeated during cycling are shown in bold.

| ***Gene (bp amplified)*** | ***Primers (5' - 3')*** | ***PCR conditions*** | ***PCR reagents*** |
|---|---|---|---|
| 16S rRNA gene, V4 region<br>254 bp<br>(Caporaso et al., 2011) | 515F:<br>GTGCCAGCMGCCGCGGTAA<br>806R:<br>GGACTACHVGGGTWTCTAAT | 94 °C for 3 min<br>**94 °C for 45 s**<br>**50 °C for 60 s**<br>**72 °C for 90 s**<br>72 °C for 10 min | 12.5 µl KAPA PCR buffer<br>0.1 µl KAPA 3G enzyme (Roche, USA)<br>0.75 µl each primer at 10 µM<br>1.5 µl BSA at 1.5 mg/ml<br>1 µl DNA template<br>8.4 µl water |
| *nifH*<br>360 bp<br>(Poly et al., 2001) | polF:<br>TGCGAYCCSAARGCBGACTC<br>polR:<br>ATSGCCATCATYTCRCCGGA | 94 °C for 3 min<br>**94 °C for 30 s**<br>**59 °C for 60 s**<br>**72 °C for 60 s**<br>72 °C for 7 min | 5 µl 10X PCR buffer<br>4 µl of dNTPs at 2.5 mM<br>0.5 µl each primer at 20 µM<br>0.25 µl Taq Polymerase, native (ThermoFisher Scientific, USA)<br>4 µl DNA template<br>0.5 µl BSA at 1.5 mg/ml<br>35.25 µl water |
| *soxB*<br>473 bp<br>(Petri et al., 2001) | 693F:<br>ATCGGNCARGCNTTYCCNTA<br>1164B:<br>AARTTNCCNCGNCGRTA | 94 °C for 3 min<br>**94 °C for 30 s**<br>**53 °C for 60 s**<br>**72 °C for 60 s**<br>72 °C for 7 min | |
| *aprA*<br>400 bp | aprA-1-FW:<br>TGGCAGATCATGATYMAYGG | 94 °C for 3 min<br>**94 °C for 30 s**<br>**53 °C for 60 s** | |

| | | |
|---|---|---|
| (Aoki et al., 2015) | aprA-5-RV: GCGCCAACNGGDCCRTA | **72 °C for 60 s**<br>72 °C for 7 min |
| *dsrA*<br>327 bp<br>(Gao et al., 2022) | DSR1F: ACSCACTGGAAGCACGGCGG | 94 °C for 3 min<br>**94 °C for 30 s**<br>**59 °C for 60 s** |
| | DSR-R: GTGGMRCCGTGCAKRTTGG | **72 °C for 60 s**<br>72 °C for 7 min |

**Table S2. PERMANOVA results for taxonomic and functional communities present across sample types.** An initial PERMANOVA is performed to determine the overall influence of sample type on the community structure. Further pairwise PERMANOVA analysis is performed to determine the compositionality differences between pairs of sample types, except for the archaeal data, where only rhizoplane and sediment are compared. All samples are retained for this analysis. Statistically significant results are indicated in bold text ($p \leq 0.05$).

| ***Gene*** | ***p-value*** | ***Pairs different*** |
|---|---|---|
| 16S rDNA (bacteria) | **0.001** | |
| | **0.001** | **phyllosphere - rhizoplane** |
| | **0.001** | **phyllosphere - sediment** |
| | **0.001** | **rhizoplane - sediment** |
| 16S rDNA (archaea) | **0.001** | |
| *nifH* | **0.001** | |
| | **0.001** | **phyllosphere - rhizoplane** |
| | **0.001** | **phyllosphere - sediment** |
| | **0.001** | **rhizoplane - sediment** |
| *soxB* | **0.001** | |
| | **0.001** | **phyllosphere - rhizoplane** |
| | **0.001** | **phyllosphere - sediment** |
| | **0.001** | **rhizoplane - sediment** |
| *aprA* | **0.001** | |
| | **0.001** | **phyllosphere - rhizoplane** |
| | **0.001** | **phyllosphere - sediment** |
| | **0.001** | **rhizoplane - sediment** |
| *dsrA* | **0.001** | |
| | **0.001** | **phyllosphere - rhizoplane** |
| | **0.001** | **phyllosphere - sediment** |
| | **0.001** | **rhizoplane - sediment** |

**Table S3. Nested LMER model ANOVA and post hoc pairwise estimated marginal means (emmeans) results for the diversity of sediment communities across succession stages.** Chao1 richness and Pielou's evenness are compared between sediment at pre-succession, early and late stages using a model that nests quadrats within patches. The Bonferroni correction is applied to pairwise emmeans p-value results. Statistically significant results are indicated in bold text ($p \leq 0.05$).

| ***Gene*** | ***Diversity index*** | ***p-value*** | ***Pairs different*** |
|---|---|---|---|
| 16S rDNA (bacteria) | Chao1 richness | **0.0022** | |
| | | **0.0024** | **early-late** |
| | | 0.2193 | pre-late |
| | | 0.6269 | pre-early |
| | Evenness | 0.1713 | |
| | | 0.2649 | early-late |
| | | 1.0000 | pre-late |
| | | 0.5851 | pre-early |
| 16S rDNA (archaea) | Chao1 richness | 0.3662 | |
| | | 0.5790 | early-late |
| | | 1.0000 | pre-late |
| | | 1.0000 | pre-early |
| | Evenness | 0.5506 | |
| | | 1.0000 | early-late |
| | | 1.0000 | pre-late |
| | | 0.8679 | pre-early |
| *nifH* | Chao1 richness | **0.0108** | |
| | | **0.0122** | **early-late** |
| | | 0.2949 | pre-late |
| | | 1.0000 | pre-early |
| | Evenness | 0.8615 | |
| | | 1.0000 | early-late |
| | | 1.0000 | pre-late |
| | | 1.0000 | pre-early |
| *soxB* | Chao1 richness | 0.3065 | |
| | | 0.7644 | early-late |
| | | 0.5388 | pre-late |
| | | 1.0000 | pre-early |
| | Evenness | 0.1816 | |
| | | 0.2425 | early-late |
| | | 0.7789 | pre-late |
| | | 1.0000 | pre-early |
| *aprA* | Chao1 richness | 0.3998 | |
| | | 0.5645 | early-late |
| | | 1.0000 | pre-late |
| | | 1.0000 | pre-early |
| | Evenness | 0.7358 | |
| | | 1.0000 | early-late |
| | | 1.0000 | pre-late |

| | | | |
|---|---|---|---|
| | | 1.0000 | pre-early |
| *dsrA* | Chao1 richness | 0.8375 | |
| | | 1.0000 | early-late |
| | | 1.0000 | pre-late |
| | | 1.0000 | pre-early |
| | Evenness | 0.2098 | |
| | | 0.4373 | early-late |
| | | 1.0000 | pre-late |
| | | 0.4404 | pre-early |

**Table S4. PERMANOVA results for sediment community structure comparisons across succession stages.** An initial PERMANOVA is performed to determine the overall influence of succession stage on the community structure. Further pairwise PERMANOVA analysis is performed to determine the compositionality differences between pairs of succession stages. Statistically significant results are indicated in bold text ($p \leq 0.05$).

| ***Gene*** | ***p-value*** | ***Pairs different*** |
|---|---|---|
| 16S rDNA (bacteria) | **0.031** | |
| | **0.008** | **early-late** |
| | 0.250 | pre-late |
| | 0.375 | pre-early |
| 16S rDNA (archaea) | **0.035** | |
| | **0.050** | **early-late** |
| | 0.125 | pre-late |
| | 0.375 | pre-early |
| *nifH* | **0.001** | |
| | **0.008** | **early-late** |
| | 0.063 | pre-late |
| | 0.250 | pre-early |
| *soxB* | 0.154 | |
| | 0.074 | early-late |
| | 0.563 | pre-late |
| | 0.125 | pre-early |
| *aprA* | **0.003** | |
| | **0.002** | **early-late** |
| | 0.188 | pre-late |
| | 0.313 | pre-early |
| *dsrA* | **0.002** | |
| | **0.008** | **early-late** |
| | 0.063 | pre-late |
| | 0.125 | pre-early |

**Table S5. Linear model analysis results for the relationship between diversity indices and plant distance from the patch edge.** Distance of plant growth from the patch edge is used as a proxy for succession, being used to test whether diversity indices increase with succession. Statistically significant results are indicated in bold text ($p \leq 0.05$).

| ***Gene*** | ***Diversity index*** | ***p-value*** | ***$R^2$*** | ***Model formula*** |
|---|---|---|---|---|
| 16S rDNA (bacteria) | Chao1 richness | **0.00** | **0.6318** | **y = 42.1x + 909** |
| | Evenness | 0.16 | 0.1105 | y = 0.002x + 0.913 |
| 16S rDNA (archaea) | Chao1 richness | 0.33 | 0.0065 | y = 3.44x + 38.2 |
| | Evenness | 0.56 | -0.0607 | y = 0.002x + 0.880 |
| *nifH* | Chao1 richness | **0.05** | **0.2622** | **y = 47.6x + 738** |
| | Evenness | 0.86 | -0.0962 | y = 0.001x + 0.878 |
| *soxB* | Chao1 richness | 0.29 | 0.0224 | y = 25.7x + 1376 |
| | Evenness | 0.15 | 0.1151 | y = 0.003x + 0.894 |
| *aprA* | Chao1 richness | 0.52 | -0.0540 | y = 26.0x + 1018 |
| | Evenness | 0.41 | -0.0231 | y = -0.002x + 0.915 |
| *dsrA* | Chao1 richness | 0.27 | 0.0317 | y = 31.8x + 1012 |
| | Evenness | 0.27 | 0.0300 | y = 0.004x + 0.868 |

**Table S6. Kruskal-Wallis and paired post hoc pairwise Dunn's Test results for the 16S rDNA marker gene (bacteria).** Comparisons of Chao1 richness and Pielou's evenness indices are made between sample types at each succession stage. The default Bonferroni correction method is used for calculating adjusted p-values for Dunn's Test results. Statistically significant results are indicated in bold text ($p \leq 0.05$).

| ***Diversity index*** | ***Succession stage*** | ***Test*** | ***p-value*** | ***Pairs different*** |
|---|---|---|---|---|
| Chao1 richness | Early | Kruskal-Wallis | **0.0038** | |
| | | dunnTest | 1.0000 | phyllosphere - rhizoplane |
| | | | **0.0061** | **phyllosphere - sediment** |
| | | | **0.0293** | **rhizoplane - sediment** |
| | Late | Kruskal-Wallis | **0.0036** | |
| | | dunnTest | 1.0000 | phyllosphere - rhizoplane |
| | | | **0.0386** | **phyllosphere - sediment** |
| | | | **0.0043** | **rhizoplane - sediment** |
| Evenness | Early | Kruskal-Wallis | **0.0212** | |
| | | dunnTest | 0.2796 | phyllosphere - rhizoplane |
| | | | 0.7587 | phyllosphere - sediment |
| | | | **0.0169** | **rhizoplane - sediment** |
| | Late | Kruskal-Wallis | **0.0008** | |
| | | dunnTest | 0.1752 | phyllosphere - rhizoplane |
| | | | 0.1752 | phyllosphere - sediment |
| | | | **0.0005** | **rhizoplane - sediment** |

**Table S7. Kruskal-Wallis and paired post hoc pairwise Dunn's Test results for the *soxB* gene.** Comparisons of Chao1 richness and Pielou's evenness indices are made between sample types at each succession stage. The default Bonferroni correction method is used for calculating adjusted p-values for Dunn's Test results. Statistically significant results are indicated in bold text ($p \leq 0.05$).

| ***Diversity index*** | ***Succession stage*** | ***Test*** | ***p-value*** | ***Pairs different*** |
|---|---|---|---|---|
| Chao1 richness | Early | Kruskal-Wallis | **0.0306** | |
| | | dunnTest | 0.0763 | phyllosphere - rhizoplane |
| | | | 1.0000 | phyllosphere - sediment |
| | | | **0.0494** | **rhizoplane - sediment** |
| | Late | Kruskal-Wallis | **0.0029** | |
| | | dunnTest | **0.0242** | **phyllosphere - rhizoplane** |
| | | | 1.0000 | phyllosphere - sediment |
| | | | **0.0043** | **rhizoplane - sediment** |
| Evenness | Early | Kruskal-Wallis | **0.0049** | |
| | | dunnTest | **0.0047** | **phyllosphere - rhizoplane** |
| | | | 1.0000 | phyllosphere - sediment |
| | | | **0.0494** | **rhizoplane - sediment** |
| | Late | Kruskal-Wallis | **0.0029** | |
| | | dunnTest | **0.0246** | **phyllosphere - rhizoplane** |
| | | | 1.0000 | phyllosphere - sediment |
| | | | **0.0043** | **rhizoplane - sediment** |

**Table S8. Kruskal-Wallis results for the 16S rDNA marker gene (archaea).** Comparisons of Chao1 richness and Pielou's evenness indices are made between sediment and rhizoplane at each succession stage. Statistically significant results are indicated in bold text ($p \leq 0.05$).

| ***Diversity index*** | ***Succession stage*** | ***Test*** | ***p-value*** |
|---|---|---|---|
| Chao1 richness | Early | Kruskal-Wallis | 0.0547 |
| | Late | Kruskal-Wallis | **0.0357** |
| Evenness | Early | Kruskal-Wallis | 0.7150 |
| | Late | Kruskal-Wallis | 1.0000 |

**Table S9. Kruskal-Wallis and paired post hoc pairwise Dunn's Test results for the *nifH* gene.** Comparisons of Chao1 richness and Pielou's evenness indices are made between sample types at each succession stage. The default Bonferroni correction method is used for calculating adjusted p-values for Dunn's Test results. Statistically significant results are indicated in bold text ($p \leq 0.05$).

| ***Diversity index*** | ***Succession stage*** | ***Test*** | ***p-value*** | ***Pairs different*** |
|---|---|---|---|---|
| Chao1 richness | Early | Kruskal-Wallis | **0.0015** | |
| | | dunnTest | **0.0009** | **phyllosphere - rhizoplane** |
| | | | 0.2590 | phyllosphere - sediment |
| | | | 0.1492 | rhizoplane - sediment |
| | Late | Kruskal-Wallis | **0.0011** | |
| | | dunnTest | 0.0916 | phyllosphere - rhizoplane |
| | | | 0.3900 | phyllosphere - sediment |
| | | | **0.0007** | **rhizoplane - sediment** |
| Evenness | Early | Kruskal-Wallis | **0.0097** | |
| | | dunnTest | 0.1418 | phyllosphere - rhizoplane |
| | | | 0.8322 | phyllosphere - sediment |
| | | | **0.0076** | **rhizoplane - sediment** |
| | Late | Kruskal-Wallis | **0.0021** | |
| | | dunnTest | 0.0799 | phyllosphere - rhizoplane |
| | | | 0.6408 | phyllosphere - sediment |
| | | | **0.0016** | **rhizoplane - sediment** |

**Table S10. Kruskal-Wallis and paired post hoc pairwise Dunn's Test results for the *aprA* gene.** Comparisons of Chao1 richness and Pielou's evenness indices are made between sample types at each succession stage. The default Bonferroni correction method is used for calculating adjusted p-values for Dunn's Test results. Statistically significant results are indicated in bold text ($p \leq 0.05$).

| ***Diversity index*** | ***Succession stage*** | ***Test*** | ***p-value*** | ***Pairs different*** |
|---|---|---|---|---|
| Chao1 richness | Early | Kruskal-Wallis | **0.0050** | |
| | | dunnTest | **0.0035** | **phyllosphere - rhizoplane** |
| | | | 0.5102 | phyllosphere - sediment |
| | | | 0.1570 | rhizoplane - sediment |
| | Late | Kruskal-Wallis | **0.0031** | |
| | | dunnTest | **0.0206** | **phyllosphere - rhizoplane** |
| | | | 1.0000 | phyllosphere - sediment |
| | | | **0.0051** | **rhizoplane - sediment** |
| Evenness | Early | Kruskal-Wallis | **0.0061** | |
| | | dunnTest | **0.0082** | **phyllosphere - rhizoplane** |
| | | | 1.0000 | phyllosphere - sediment |
| | | | **0.0312** | **rhizoplane - sediment** |
| | Late | Kruskal-Wallis | **0.0097** | |
| | | dunnTest | 0.0602 | phyllosphere - rhizoplane |
| | | | 1.0000 | phyllosphere - sediment |
| | | | **0.0125** | **rhizoplane - sediment** |

**Table S11. Kruskal-Wallis and paired post hoc pairwise Dunn's Test results for the *dsrA* gene.** Comparisons of Chao1 richness and Pielou's evenness indices are made between sample types at each succession stage. The default Bonferroni correction method is used for calculating adjusted p-values for Dunn's Test results. Statistically significant results are indicated in bold text ($p \leq 0.05$).

| ***Diversity index*** | ***Succession stage*** | ***Test*** | ***p-value*** | ***Pairs different*** |
|---|---|---|---|---|
| Chao1 richness | Early | Kruskal-Wallis | **0.0019** | |
| | | dunnTest | **0.0013** | **phyllosphere - rhizoplane** |
| | | | 0.1558 | phyllosphere - sediment |
| | | | 0.2860 | rhizoplane - sediment |
| | Late | Kruskal-Wallis | 0.0600 | |
| | | dunnTest | 0.3505 | phyllosphere - rhizoplane |
| | | | 1.0000 | phyllosphere - sediment |
| | | | 0.0602 | rhizoplane - sediment |
| Evenness | Early | Kruskal-Wallis | **0.0013** | |
| | | dunnTest | **0.0008** | **phyllosphere - rhizoplane** |
| | | | 0.2021 | phyllosphere - sediment |
| | | | 0.1693 | rhizoplane - sediment |
| | Late | Kruskal-Wallis | **0.0083** | |
| | | dunnTest | 0.5831 | phyllosphere - rhizoplane |
| | | | 0.2231 | phyllosphere - sediment |
| | | | **0.0062** | **rhizoplane - sediment** |

**Table S12. Nested LMER model ANOVA results for the diversity of rhizoplane communities across succession stages.** Chao1 richness and Pielou's evenness are compared between rhizoplane communities at early and late stages using a model that nests quadrats within patches. Statistically significant results are indicated in bold text ($p \leq 0.05$).

| ***Gene*** | ***Diversity index*** | ***p-value*** |
|---|---|---|
| 16S rDNA (bacteria) | Chao1 richness | 0.5237 |
| | Evenness | 0.9703 |
| 16S rDNA (archaea) | Chao1 richness | 0.1120 |
| | Evenness | 0.8530 |
| *nifH* | Chao1 richness | 0.6345 |
| | Evenness | 0.4507 |
| *soxB* | Chao1 richness | 0.1947 |
| | Evenness | **0.0245** |
| *aprA* | Chao1 richness | 0.2750 |
| | Evenness | 0.5520 |
| *dsrA* | Chao1 richness | 0.6581 |
| | Evenness | 0.0786 |

**Table S13. Nested LMER model ANOVA results for the diversity of phyllosphere communities across succession stages.** Chao1 richness and Pielou's evenness are compared between phyllosphere communities at early and late stages using a model that nests quadrats within patches. Statistically significant results are indicated in bold text ($p \leq 0.05$).

| ***Gene*** | ***Diversity index*** | ***p-value*** |
|---|---|---|
| 16S rDNA (bacteria) | Chao1 richness | 0.0885 |
| | Evenness | 0.7055 |
| *nifH* | Chao1 richness | 0.2007 |
| | Evenness | 0.9457 |
| *soxB* | Chao1 richness | 0.6247 |
| | Evenness | 0.8523 |
| *aprA* | Chao1 richness | 0.1964 |
| | Evenness | 0.1718 |
| *dsrA* | Chao1 richness | **0.0023** |
| | Evenness | **0.0012** |

**Table S14. PERMANOVA results for rhizoplane community structure comparisons across succession stages.** Community composition is compared between rhizoplane communities at early and late succession stages. Statistically significant results are indicated in bold text ($p \leq 0.05$).

| ***Gene*** | ***p-value*** |
|---|---|
| 16S rDNA (bacteria) | **0.010** |
| 16S rDNA (archaea) | **0.022** |
| *nifH* | **0.010** |
| *soxB* | **0.010** |
| *aprA* | **0.010** |
| *dsrA* | **0.010** |

**Table S15. PERMANOVA results for phyllosphere community structure comparisons across succession stages.** Community composition is compared between phyllosphere communities at early and late succession stages. Statistically significant results are indicated in bold text ($p \leq 0.05$).

| ***Gene*** | ***p-value*** |
|---|---|
| 16S rDNA (bacteria) | **0.017** |
| *nifH* | **0.023** |
| *soxB* | **0.031** |
| *aprA* | **0.016** |
| *dsrA* | **0.010** |

### SUPPLEMENTARY FIGURES

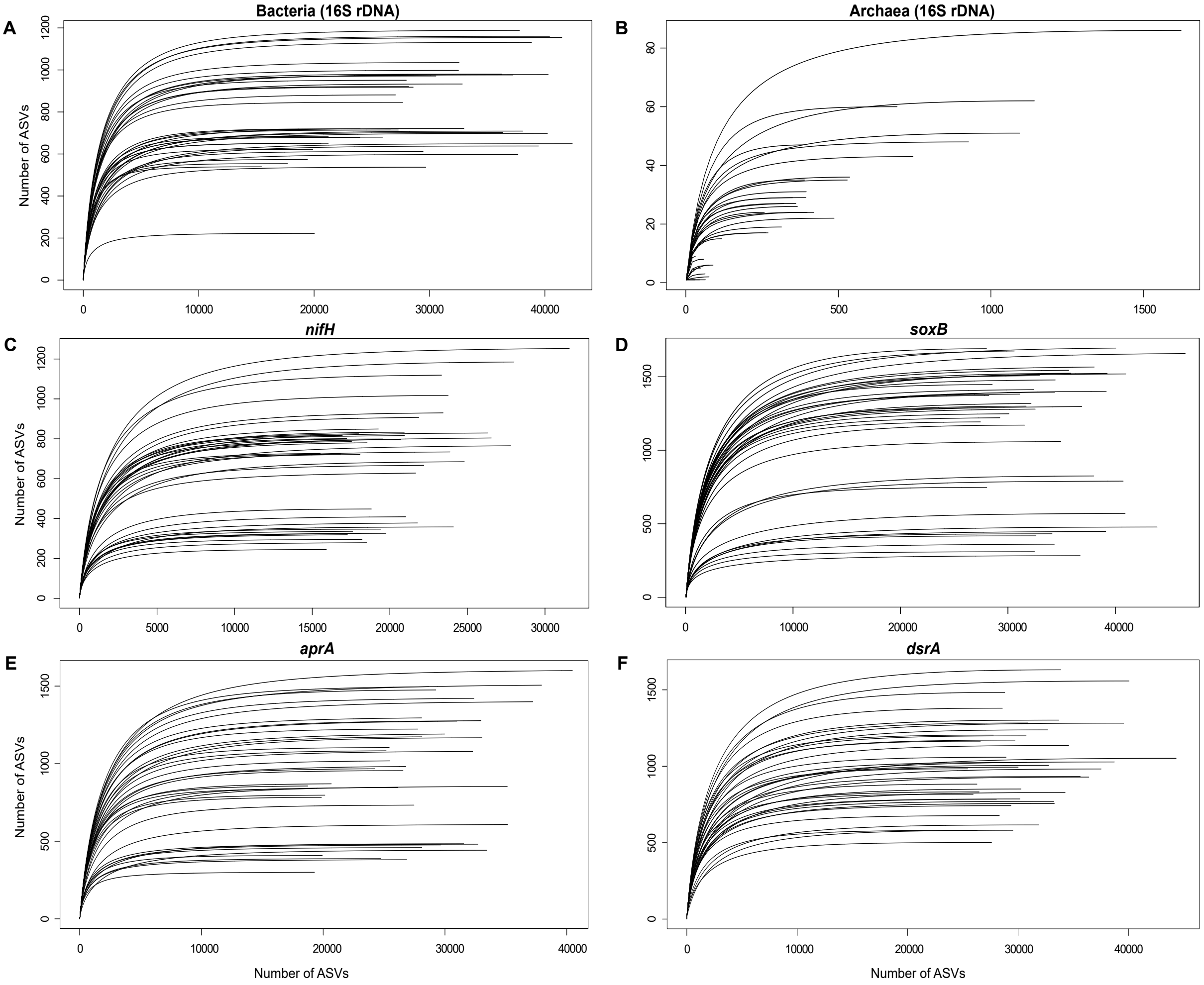


**Fig S1. Rarefaction curves for the datasets analysed.** Curves for the bacterial (16S rDNA marker) **(A)**, archaeal (16S rDNA marker) **(B)**, *nifH* **(C)**, *soxB* **(D)**, *aprA* **(E)** and *dsrA* **(F)** datasets. Sufficient sequencing depth is achieved for all the amplified genes, as evidenced by the plateauing number of ASVs, except for the archaeal dataset, which was subset from the 16S rDNA amplicon data.

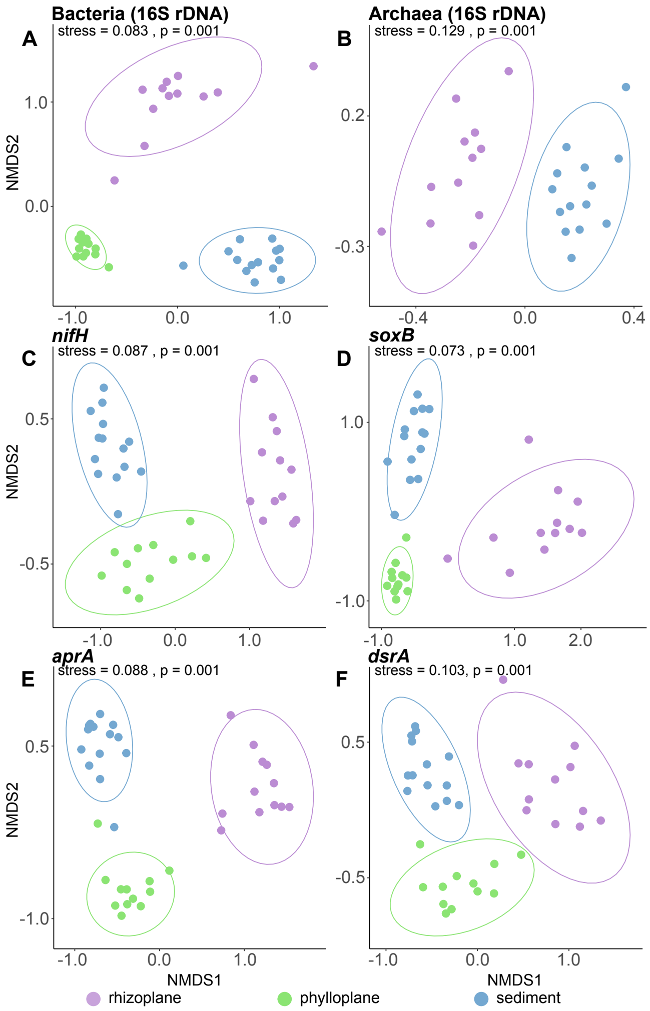


**Fig S2. NMDS ordinations of entire taxonomic and functional communities present in different sample types.** Composition is different between sample types (PERMANOVA, $p \leq 0.05$), for bacterial **(A)**, archaeal **(B)**, *nifH* **(C)**, *soxB* **(D)**, *aprA* **(E)**, and *dsrA* **(F)** communities in the rhizoplane, phyllosphere and sediment. All samples are retained for these ordination plots. Colours represent the sample types.

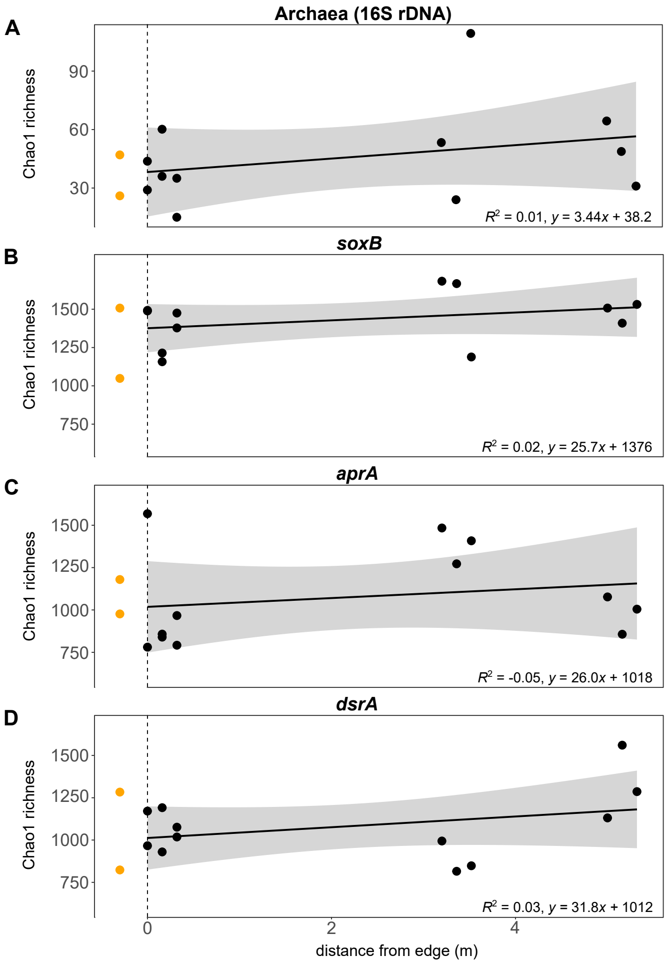


**Fig S3. The distance-diversity relationship for the Chao1 richness index, for sediment communities.** Richness does not increase with succession for the archaeal **(A)**, *soxB* **(B)**, *aprA* **(C)**, and *dsrA* **(D)** communities in the sediment. Distance is used here as a substitute for time, following the chronosequence used in this study. Pre-succession sediment sampled outside the seagrass patch is indicated by the orange points. Grey-shaded areas in the plot indicate the confidence interval (95%) of the best fit line.

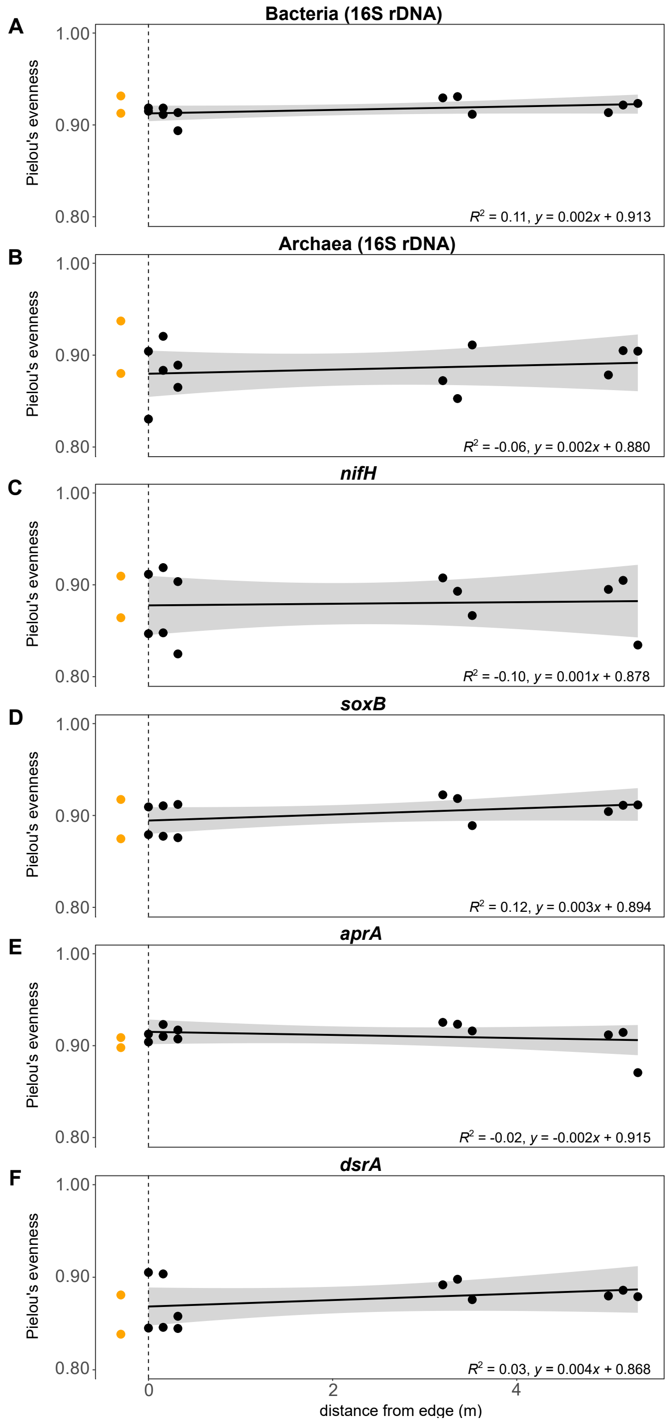


**Fig S4. The distance-diversity relationship for Pielou's evenness index, for sediment communities.** Evenness does not increase with succession for any of the communities investigated, which include, the bacterial **(A)**, archaeal **(B)**, nifH **(C)**, soxB **(D)**, aprA **(E)**, and dsrA **(F)** communities. Distance is used here as a substitute for time, following the chronosequence used in this study. Pre-succession sediment sampled outside the seagrass patch is indicated by the orange points. Grey-shaded areas in the plot indicate the confidence interval (95%) of the best fit line.

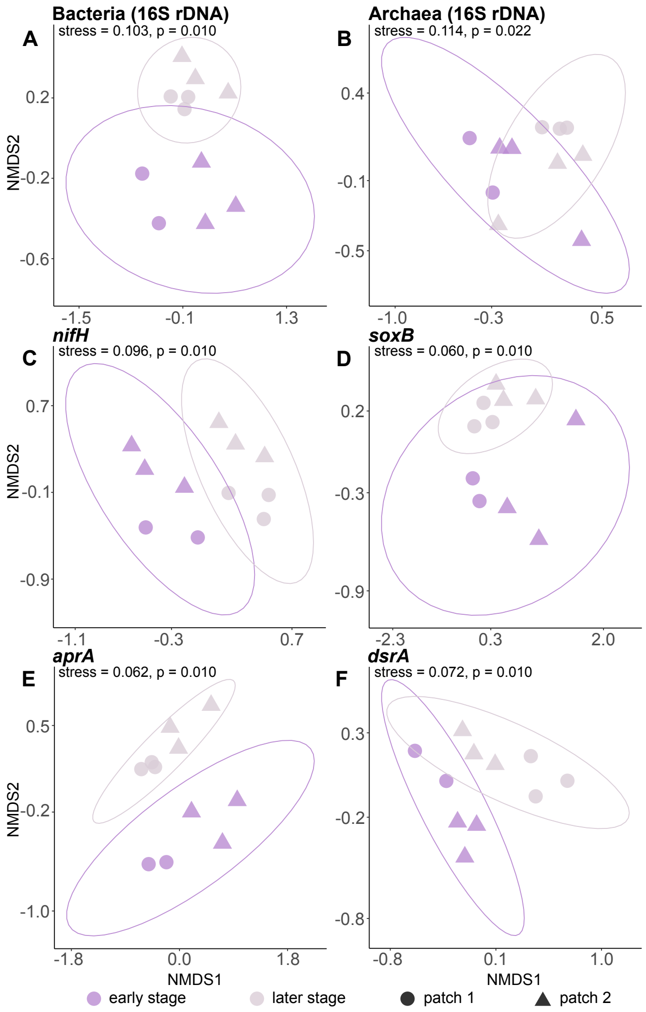


**Fig S5. NMDS ordinations of rhizoplane communities between chronosequence stages.** Community composition varies between the early and late stages of the chronosequence (PERMANOVA, $p \leq 0.05$), for bacterial **(A)**, archaeal **(B)**, *nifH* **(C)**, *soxB* **(D)**, *aprA* **(E)**, and *dsrA* **(F)** communities in the rhizoplane. A rhizoplane sample from the edge of patch 1 (darker coloured circle) has been removed from the above ordination plots and related statistical testing. Colours represent the stages across the chronosequence, while shapes represent the replicate patches sampled.

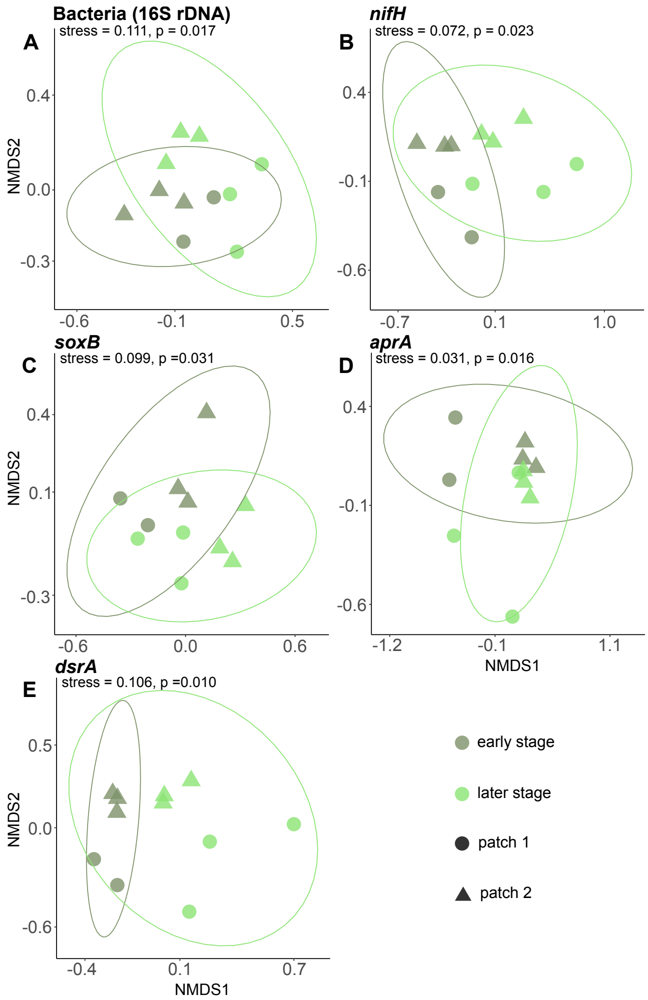


**Fig S6. NMDS ordinations of phyllosphere communities between chronosequence stages.** Community composition varies between the early and late stages of the chronosequence (PERMANOVA, $p \leq 0.05$), for bacteria **(A)**, *nifH* **(B)**, *soxB* **(C)**, *aprA* **(D)**, and *dsrA* **(E)** communities in the phyllosphere. A phyllosphere sample from the edge of patch 1 (darker coloured circle) has been removed from the above ordination plots and related statistical testing. Colours represent the stages across the chronosequence, while shapes represent the replicate patches sampled.

**Supplementary materials references**

Aoki, M., Kakiuchi, R., Yamaguchi, T., Takai, K., Inagaki, F., & Imachi, H. (2015). Phylogenetic diversity of *aprA* genes in subseafloor sediments on the Northwestern Pacific margin off Japan. *Microbes Environ*, *30*(3), 276-280. https://doi.org/10.1264/jsme2.ME15023

Caporaso, J. G., Lauber, C. L., Walters, W. A., Berg-Lyons, D., Lozupone, C. A., Turnbaugh, P. J., Fierer, N., & Knight, R. (2011). Global patterns of 16S rRNA diversity at a depth of millions of sequences per sample. *Proc Natl Acad Sci U S A*, *108 Suppl 1*(Suppl 1), 4516-4522. https://doi.org/10.1073/pnas.1000080107

Gao, H., Wang, C., Chen, J., Wang, P., Zhang, J., Zhang, B., Wang, R., & Wu, C. (2022). Enhancement effects of decabromodiphenyl ether on microbial sulfate reduction in eutrophic lake sediments: A study on sulfate-reducing bacteria using *dsrA* and *dsrB* amplicon sequencing. *Science of The Total Environment*, *843*, 157073. https://doi.org/https://doi.org/10.1016/j.scitotenv.2022.157073

Petri, R., Podgorsek, L., & Imhoff, J. F. (2001). Phylogeny and distribution of the *soxB* gene among thiosulfate-oxidizing bacteria. *FEMS Microbiol Lett*, *197*(2), 171-178. https://doi.org/10.1111/j.1574-6968.2001.tb10600.x

Poly, F., Monrozier, L. J., & Bally, R. (2001). Improvement in the RFLP procedure for studying the diversity of *nifH* genes in communities of nitrogen fixers in soil. *Res Microbiol*, *152*(1), 95-103. https://doi.org/10.1016/s0923-2508(00)01172-4
